# SERRATE interacts with the nuclear exosome targeting (NEXT) complex to degrade primary miRNA precursors in Arabidopsis

**DOI:** 10.1101/2020.04.08.032003

**Authors:** Mateusz Bajczyk, Heike Lange, Dawid Bielewicz, Lukasz Szewc, Susheel Sagar Bhat, Jakub Dolata, Lauriane Kuhn, Zofia Szweykowska-Kulinska, Dominique Gagliardi, Artur Jarmolowski

**Author notes:** correspondence should be addressed to: Heike Lange, Artur Jarmolowski.

## Abstract

SERRATE/ARS2 is a conserved RNA effector protein involved in transcription, processing and export of different types of RNAs. In Arabidopsis, the best-studied function of SERRATE (SE) is to promote miRNA processing. Here, we report that SE interacts with the Nuclear Exosome Targeting (NEXT) complex, comprising the RNA helicase HEN2, the RNA binding protein RBM7 and one of the two zinc-knuckle proteins ZCCHC8A/ZCCHC8B. The identification of common targets of SE and HEN2 by RNA-seq supports the idea that SE cooperates with NEXT for RNA surveillance by the nuclear exosome. Among the RNA targets accumulating in absence of SE or NEXT are miRNA precursors. Loss of NEXT components results in the accumulation of pri-miRNAs without affecting levels of miRNAs, indicating that NEXT is, unlike SE, not required for miRNA processing. As compared to *se-2*, *se-2 hen2-2* double mutants showed increased accumulation of pri-miRNAs, but partially restored levels of mature miRNAs and attenuated developmental defects. We propose that the slow degradation of pri-miRNAs caused by loss of HEN2 compensates for the poor miRNA processing efficiency in *se-2* mutants, and that SE regulates miRNA biogenesis through its double contribution in promoting miRNA processing but also pri-miRNA degradation through the recruitment of the NEXT complex.

## Introduction

In plants, many miRNAs are encoded by independent RNA polymerase II transcription units. The primary transcripts contain a 5′ cap structure as well as a poly(A) tail at the 3′ end (1), and sometimes introns. The primary or spliced pri-miRNAs adopt stem loop structures which are processed by the nuclear endonuclease Dicer like 1 (DCL1) (2) in two sequential reactions. The first step creates shorter miRNA precursors called pre-miRNAs. The second step produces miRNA/miRNA* duplexes of mostly 21 nt with 2 nt overhangs at both 3′ ends. DCL1 associates with the double stranded RNA binding protein HYPONASTIC LEAVES 1 (HYL1) and the zinc finger domain-containing RNA effector protein SERRATE (3–5). Both HYL1 and SE interact with DCL1 as well as with each other (6–8). DCL1, SE and HYL1 form the core of the plant Microprocessor complex and co-localize within the nucleus in special structures known as dicing bodies (D-bodies) (7). In absence of HYL1 or upon reduced activity of SE, dicing by DCL1 is not completely abolished but less efficient and less accurate (3–5). The importance of SE for the biogenesis of miRNAs is illustrated by the phenotype of *se* mutants. Complete loss of SE expression as in the null mutant *se-4* is embryonic lethal, while the *se-1* mutation, which results in the expression of a truncated SE lacking 20 amino acids at its C-terminus, severely affects developmental timing, phylotaxy, meristem function and patterning of leaves and flowers (4,9). The *se-2* mutation, in which the SE protein is truncated by 40 amino acids from the C-terminus, additionally displays the hyponastic leave shape that is characteristic for many miRNA biogenesis mutants including *hyl1-2, hst-1 and ago1-25* (10–13).

In addition to its role in miRNA biogenesis, Arabidopsis SE contributes to both constitutive and alternative splicing of pre-mRNAs (14,15), enhances the transcription of intronless genes (16), and regulates the expression of transposons (17). The human SE ortholog, ARSENITE RESISTANCE PROTEIN 2 (ARS2), is also required for miRNA biogenesis (18). In addition, ARS2 participates to the transcription termination and degradation of 3′ extended snRNAs, PROMPTs (promoter upstream transcripts), eRNAs (enhancer RNAs) and RNAs with 3′ ends inside first introns of protein coding genes (18–21) as well as to the 3′ end processing of histone mRNAs (22) and nuclear export of both snRNAs and mRNAs (23). The roles of SE/ARS2 as versatile and multifunctional RNA effector proteins are linked to their capability to associate with various RNA-protein complexes (20,21,24). An important partner of both human ARS2 and Arabidopsis SE is the nuclear cap-binding complex (CBC) which binds to the 5′ cap structure of all nascent RNA polymerase II transcripts (15,18,25). For example, human CBC-ARS2 interacts with Drosha for miRNA processing, with PHAX (phosphorylated adapter for RNA export) for snRNA export, or with ALYREF, a subunit of the THO/TREX complex that couples mRNA transcription to export (23). Alternatively, CBC-ARS2 can associate with the poly(A) tail exosome targeting (PAXT) and the nuclear exosome targeting (NEXT) complexes to promote RNA degradation by the RNA exosome (21,26,27). PAXT and NEXT are so-called activator-adapter complexes that assist the RNA exosome in the nucleoplasm, while distinct activator-adapters such as the TRAMP and SKI complexes modulate the activity of the RNA exosome in the nucleolus and cytoplasm, respectively (28).

PAXT connects the RNA helicase hMTR4 and the Zn-knuckle protein ZFC3H1 with the nuclear poly(A)-binding protein PABPN1, and promotes the degradation of transcripts with long poly(A) tails in the nucleoplasm. In *Schizosaccharomyces pombe*, homologs of MTR4, ZFC3H1 and PAPBN1 form a similar complex named MTREC (Mtl1-Red1-Core) or NURS (nuclear RNA silencing) which is required for the elimination of meiotic mRNAs during mitosis and the degradation of unspliced mRNAs, non-coding RNAs, and cryptic intergenic transcripts (29,30). The human NEXT complex also comprises hMTR4 but in association with the Zn-knuckle protein ZCCHC8 and the RNA binding protein RBM7, and targets snRNA, PROMPTs, eRNA, and histone RNA precursors extended at 3′ ends for degradation by the nucleoplasmic RNA exosome (31–33). Both PAXT and NEXT assemble in higher order complexes with the Zn-finger protein ZC3H18 and CBC-ARS2 (26). The current view is that the mutually exclusive interaction of CBC-ARS with ZC3H18/NEXT/PAXT or with PHAX/ALYREF determines whether a nuclear RNA is degraded or exported to the cytoplasm (23,24,34).

In a previous study, we demonstrated that the Arabidopsis RNA exosome complex is associated with two nuclear RNA helicases, AtMTR4 and HEN2 (35). AtMTR4 is restricted to nucleoli and targets predominantly ribosomal RNA precursors and rRNA maturation by-products (36). HEN2 is located in the nucleoplasm and targets diverse RNAs including mis-spliced mRNAs, excised introns, 3′ extended snoRNAs and intergenic transcripts (35). Co-purification experiments identified two Zn-knuckle proteins, ZCCHC8A and ZCCHC8B, and a homolog of the RNA binding protein RBM7 as components of the Arabidopsis NEXT complex. This experiment also co-purified CBP20 and CBP80, the two subunits of the cap-binding complex. Whether SE can also associate with the plant NEXT complex was not studied yet.

Here, we show that Arabidopsis SE is associated with both CBC and NEXT *in vivo*, and investigate the individual interactions between SE and subunits of NEXT. We show by yeast two hybrid assays, FRET-FLIM and reciprocal co-immunoprecipitation experiments that the two ZCCHC8A and ZCCHC8B subunits directly interact with SE and also function as the central scaffold of NEXT. Loss of HEN2, RBM7, and simultaneous loss of both ZCCHC8A and ZCCHC8B results in the accumulation of miRNA precursors and miRNA maturation by-products, indicating that NEXT stimulates their degradation by the RNA exosome. Further support for a cooperation of SE and the NEXT complex in RNA surveillance is provided by the identification of common targets of SE and HEN2 by transcriptome-wide analyses. Interestingly, we observe that the accumulation of primary miRNAs caused by loss of HEN2 restores levels of mature miRNAs and mostly rescues the developmental phenotype of *se-2* mutants. Taken together, our data suggest that SE is not only required for miRNA processing but also actively promotes the degradation of pri-miRNAs and other types of RNAs by recruiting the nuclear exosome targeting complex. We propose that SE fine-tunes miRNA biogenesis by facilitating miRNA processing or recruiting the NEXT complex to trigger the degradation of pri-miRNAs.

## MATERIALS AND METHODS

### Plant material

All plants used in this study are of Columbia accession. If not noted otherwise, plants were grown on soil at 22 °C with 16 h light and 8 h darkness. The *hen2-2* T-DNA insertion line has been previously described (35). The viable exosome mutant lines *mtr3* and *rrp4KD* lines have been described in (37,38). Other T-DNA insertion Arabidopsis lines were obtained from the NASC Arabidopsis mutant collection (http://arabidopsis.info/): *zcchc8a-1* (SAIL_1230_H09), *zcchc8a-2* GK_598CO6, *zcchc8b-1* (GK_667A12), *rbm7-1* (SALK_005077), *se-2* (SAIL_44_G12), *hyl1-2* (SALK_064863). The double mutants *zcchc8a-1 zcchc8b-1* (*zch8a zch8b), hyl1-2 hen2-2, and se-2 hen2-2* were obtained by crosses. The transgenic line expressing FLAG-SERRATE under the control of 35S CaMV promoter in the *se-1* mutant background has been described previously (15).

### Expression vectors

For co-immunoprecipitation experiments, the genomic sequences of *ZCCHC8A* (*AT5G38600*) and *RBM7* (*AT4G10110*) including their putative promoter regions of 1600 bp and 1200 bp upstream of the transcription start sites, respectively, but lacking the stop codon and the 3′ UTR were cloned into vector pGWB604 (39). The genomic sequence of *ZCCHC8B* was cloned without its putative promoter but with the 5′ UTR into vector Ub10:CGFP (40). Transgenic lines were obtained by Agrobacterium-mediated floral dip transformation (41) of *zcchc8a-2*, *rbm7-1* and Col-0 plants, respectively.

For FRET-FLIM experiments, the coding sequences of *SE, HEN2, ZCCHC8A, ZCCHC8B, RBM7 and PRP39A* were amplified from cDNA and cloned into a modified pSAT6-DEST-EGFP vector (42), in which the 35S CaMV promoter was replaced by the *UBQ10* promoter. For RFP-fusion proteins, the sequence encoding EGFP was replaced by tagRFP. For BiFC, HEN2 was fused to the N-terminal part of Venus (nVEN-HEN2) and ZCCHC8A was fused to the C-terminal part of CFP (ZCCHC8A-cCFP) derived from vectors described in (43).

For yeast two hybrid experiments, the coding sequences of *SE*, *HEN2*, *ZCCHC8A*, *ZCCHC8B*, *RBM7* were cloned into vectors pGBKT7 and pGADT7 (Clontech) encoding the Gal4 DNA binding and Gal4 activating domain, respectively. The vectors encoding MBP-SE and MBP-GFP have been described in (11). The sequence of all primers is provided in Supplementary Table 1.

### Co-immunoprecipitation and mass-spectrometry analyses

For each co-immunoprecipitation (co-IP), 0.3 g of flowers pooled from 5-10 plants grown at 20 °C were ground in liquid nitrogen. The powder was extracted for 30 min in 1.5 ml of lysis buffer (50 mM Tris HCl pH 8.0, 50 mM NaCl, 1 % Triton X-100 and 1x cOmplete™ EDTA-free protease inhibitor (Roche)). After removal of cell debris by centrifugation (twice 10 min, 16000g, 4 °C) the cleared supernatants were incubated for 30 min with anti-GFP or anti-FLAG antibodies coupled to magnetic microbeads (μMACS GFP and DYKDDDDK isolation Kits, Miltenyi). Beads were loaded on magnetized MACS separation columns equilibrated with lysis buffer and washed four times with 300 µl washing buffer (50 mM Tris HCl pH 7.5, 0.1 % Triton X-100). Samples were eluted in 100 µl of pre-warmed elution buffer (Milteny). Control IPs were performed in Col-0 using either GFP or FLAG antibodies.

Eluted proteins were digested with sequencing-grade trypsin (Promega) and analysed by nanoLC-MS/MS on a QExactive+ mass spectrometer coupled to an EASY-nanoLC-1000 (Thermo-Fisher Scientific, USA) as described before (44). Data were searched against the TAIR10 database with a decoy strategy. Peptides were identified with Mascot algorithm (version 2.5, Matrix Science, London, UK) and the data were imported into Proline 1.4 software (http://proline.profiproteomics.fr/). The protein identification was validated using the following settings: Mascot pretty rank <= 1, FDR <=1 % for PSM scores, FDR <= 1 % for protein set scores. The total number of MS/MS fragmentation spectra was used to quantify each protein from at least two independent biological replicates. Biological replicates consisted of plants of the same genotype grown at different dates and in different growth chambers. In the case of ZCCHC8B we used plants derived from two independent transformants that were grown in parallel.

For the statistical analysis we compared the data from two independent experiments for ZCCHC8A, ZCCHC8B, RBM7, and four IPs from two biological replicates for SE to the control IPs using a homemade R package as described in (44). The R-package performs a negative-binomial test using an edgeR-based GLM regression and calculates the fold change and an adjusted p-value corrected by Benjamini–Hochberg for each identified protein. The mass spectrometry proteomics data have been deposited to the ProteomeXchange Consortium (http://proteomecentral.proteomexchange.org) via the PRIDE partner repository (45) with the dataset identifier PXD014447.

### Yeast two-hybrid analysis

The yeast two hybrid (YTH) experiments were performed in three replicates using the Matchmaker Gal4-based two-hybrid system (Clontech) and the *Saccharomyces cerevisiae* strain Y2HGold as described in (11). Yeast cells co-expressing proteins fused to the Gal4 activation domain (AD) and DNA binding domain (BD) were selected on medium lacking the amino acids leucine (L) and tryptophan (T) at 28 °C. Interaction was tested on medium lacking also histidine (H) and adenine (A), and on medium complemented with aureobasidin (Aur) and the galactosidase substrate X-αGal.

### Protoplast isolation

The epidermal layer of Arabidopsis leaves was removed using the Tape-Arabidopsis Sandwich method (46). Leaves were incubated in digestion buffer (0.4 M mannitol, 20 mM KCl, 20 mM MES pH 5.7, 1 % (w/v) Cellulase Onozuka R10 (Serva), 0.2 % (w/v) Macerozyme R10 (Serva) for 30 minutes with shaking at 20-40 rpm. Protoplasts were released by pipetting and the protoplast suspension was incubated for further 15 minutes before protoplasts were collected by centrifugation for 5 minutes at 180 x g, washed three times in buffer W5 (154 mM NaCl, 125 mM CaCl_2_, 5 mM KCl, 2 mM MES pH 5.7) and resuspended in 1 ml of 0.2 M mannitol, 15 mM MgCl_2_, 4 mM MES pH 5.7. 100 μl of protoplasts were transferred to new round-bottom tubes containing 5 μg of the desired expression vectors. After adding 100 μl PEG/Ca^2+^ solution (40 % (w/v) polyethylene glycol 4000 (PEG, Sigma-Aldrich), 0.2 M mannitol, 100 mM Ca(NO_3_)_2_) samples were incubated 5 minutes at room temperature before the transformation was stopped by diluting the sample with 450 μl of buffer W5. Protoplasts were collected by centrifugation (3 min at 300g) and resuspended in 300 μl of buffer W1 (0.5 M mannitol, 20 mM KCl, 4 mM MES pH 5.7). Protoplasts were incubated for 10-12 h at 22 °C in dark before FRET-FLIM analyses and 18-22 h at 22 °C before BiFC – FRET-FLIM experiments.

### FRET-FLIM analysis

The PicoHarp300-Dual Channel SPAD system (PicoQuant) in combination with a Nikon A1Rsi microscope armed with the 40×/1.25 water-immersion objective was used for FRET-FLIM analyses. The Picosecond Pulsed Diode Laser LDH-D-C-485 and Supercontinuum Tunable White Light Laser (488 nm) were used for generation of 100 ps excitation pulses at a repetition of 40 MHz. Images were of 256 x 256 pixels frame size and collected with an average count rate of around 10^4^ photons per second for 90-120 seconds. PhoTime 64 (PicoQuant) software was used to calculate fluorescence decay curves for each pixel. The curves were fitted with the double-exponential decay model using default parameters. For each sample, more than ten cells from at least three biological repeats (independent protoplast isolation and transformation) were used. The results were evaluated with the two-sided Mann–Whitney–Wilcoxon test (p-value < 0.001).

### Protein pull-down assay

Recombinant SE and GFP proteins fused to maltose-binding protein (MBP) were purified from *Escherichia coli* as described before (11). [^35^S]-methionine labelled HEN2, ZCCHC8A, ZCCHC8B and RBM7 proteins were obtained by *in vitro* transcription and translation using the TNT T7 coupled wheat germ extract system (Promega) following the protocol supplied by the manufacturer. Purified MBP-SE and MBP-GFP were bound to amylose beads (NEB) by overnight incubation at 4 °C. Excess protein was removed by washing the beads three times with 20 mM Tris HCl pH 7.5, 200 mM NaCl, 1 mM EDTA, cOmplete™ EDTA-free protease inhibitor (Roche) before the radiolabelled proteins were added. After incubation for 2 h at 4 °C the beads were washed five times and proteins were eluted with 20 mM maltose. Eluted proteins were separated by 12 % SDS-PAGE and detected using a FLA-5000 phosphoimager (Fujifilm).

### RNA analysis

For qRT-PCR and RNA blots, total RNA were prepared from rosettes of three biological replicates of four (Fig. 10) and five (Fig. 6) week-old plants using the Direct-zol^TM^ RNA Mini Prep Kit (Zymo Research). DNA contaminations were removed using Turbo DNase I. 4 µg RNA were used for cDNA synthesis using SuperScript III Reverse Transcriptase and oligo(dT)_18_. Real-time quantitative PCR (qPCR) was performed using Power SYBR® Green PCR Master Mix (Thermo Fisher Scientific) and the QuantStudio 7 Flex Real-Time PCR System. *GAPDH* (*AT1G13440*) and *PP2A* (*AT1G13320*) mRNAs were used for normalisation. Following log transformation, the Student’s t-test (p-value < 0.05) was used for statistical analyses. 3′ RACE-PCR was performed using the SMARTer PCR cDNA synthesis kit (Clontech) following the instructions supplied by the manufacturer. PCR products were separated by electrophoresis on a 1 % (w/v) agarose / TBE gel and visualised by ethidium bromide. PCR products amplified from the *hen2-2* samples were excised from the gel, cloned into pGEM, transformed in *E. coli* and analysed by Sanger sequencing.

### RNA-seq libraries

Total RNA was prepared from rosette leaves from three biological replicates of four week-old plants grown at 22 °C and 20 °C for the first and second RNA-seq experiments, respectively. RiboMinus™ Plant Kit for RNA-Seq (Thermo Fischer Scientific) was used to deplete rRNAs. Sequencing libraries were prepared from 300 ng of ribo-depleted RNA (NEBNext Ultra II Directional RNA Library preparation kit) and indexed with NEB Next Illumina indexing primers (NEB). An Agilent Bioanalyzer with high sensitivity DNA chip was used to confirm the quality of the libraries before they were sent for sequencing. Paired-end (2×150bp) sequencing was performed at Fasteris, Geneva, Switzerland on an HiSeq4000 platform. The first RNA-seq experiment comprised wild type and *hen2-2* with an average depth of 102 million mapped reads per library. The second RNA-seq experiment comprised wild type, *hen2-2*, *se-2* and *se-2 hen2-2* with an average depth of 39 million mapped reads per library.

### RNA-seq data analysis

Raw reads were trimmed for the first 12 nucleotides with the FASTX-Toolkit. Sequencing adapters and remaining rRNA reads were removed using Trimmomatic (47) and Bowtie (48), respectively. Cleaned reads were aligned to the Arabidopsis TAIR10 reference genome using HISAT2 (49). The overall alignment rate was 95-96 % for each sample. Stringtie and prepDE.py python scripts were used to obtain gene count matrix reads. For the Stringtie analysis, the TAIR 10.46.gtf annotation file was supplemented with pri-miRNA coordinates retrieved from the mirEX2 database (50). Boxplots were generated using R. Differential gene expression analysis was performed using the DESeq2 R package (51). For Fig. 5 and Supplementary Fig. S3, RNA-seq read distribution profiles were generated using the bedtools package (52) and visualized using Microsoft Office Excel. Plots represent mean (n=3) RPM values. *MIR* genes structures are generated based on mirEX2 (50,53). For Fig. 8, Fig. 9, Fig. S7 and Fig. S8, the cumulated reads of three replicates were visualised using IGV v 2.4.14. Venn diagrams were generated using Serial list v 2.3 and https://www.meta-chart.com/venn#/display.

### Small RNA library preparation and analysis

Total RNA was prepared from rosette leaves of 4 week-old wild type, *hen2-2, se-2, hen2-2 se-2* plants grown at 20 °C using the Direct-zol™ RNA kit (Zymo Research). RNA was quantified using the Qubit RNA Assay Kit (Life Technologies) and integrity was confirmed on the Agilent Bioanalyzer 2100 system. 10 μg of each RNA sample were separated by electrophoresis on a 15 % polyacrylamide 8M urea gel in 89 mM Tris, 89 mM borate, 2 mM EDTA. Small RNA fractions were cut out and purified from the gel. Libraries were prepared using the TruSeq Small RNA Library Preparation Kit (Illumina). Single-end (1×50bp) sequencing was performed at Fasteris, Geneva, Switzerland on a HiSeq 4000 platform. Adapter sequences were removed from raw reads with FASTX-Toolkit (fastx_clipper). Bowtie was used to align clean reads to the *Arabidopsis thaliana* TAIR10 reference genome using command ‘bowtie –v 0 –k 1 –S’ (48). To obtain mapped reads counts we used Samtools view with -h -F 4 parameters. Statistical analysis was performed with DESeq2 R package (51).

### Accession Numbers

*SERRATE: AT2G27100; ZCCHC8A: AT5G38600; ZCCHC8B: AT1G67210; RBM7: AT4G10110; HEN2: AT2G06990; HYL1: AT1G09700; PRP39A: AT1G04080*

### Data availability

The RNA-seq data are available at GEO, accession GSE131542. The proteomic data supporting the conclusions of this study are available at PRIDE PXD014447. Uncropped high-resolution pictures of plants, gels and blots, qRT-PCR data, DESeq2 data and mapping files for the visualisation of RNA-seq data are available at https://doi.org/10.6084/m9.figshare.c.4930314.v1.

## RESULTS

### SERRATE interacts with the NEXT complex

To identify the SE interactome, we performed co-immunoprecipitation experiments using an FLAG-SE fusion protein expressed in the *se-1* mutant background (15). The fusion protein restored all developmental defects of the *se-1* mutant indicating that it is a functional protein (Fig. 1A). Proteins that co-purified with FLAG-SE or in mock-IPs were identified by mass spectrometry (LC-MS/MS). Mock-IPs were performed in Col-0 using the anti-FLAG antibody. Among the proteins that were enriched in FLAG-SE IPs as compared to mock IPs (Supplementary Table 2) were the two subunits of the nuclear cap binding complex (CBC), CBP80 and CBP20 (Fig. 1B – green squares), which confirmed our previous results (15). We also detected factors that have been previously shown to interact with SE for miRNA biogenesis, namely HOS5, CPL1, DCL1, CDC5, and BRM (Fig. 1B – blue squares) (7,54,55). SE, HOS5 and CPL1 have also roles in splicing (56). CDC5 is a DNA binding protein that recruits RNA polymerase II to the promoters of miRNA genes (57). CDC5 interacts also directly with both DCL1 and SE and was therefore suggested to have a second role in miRNA processing (57). Similarly, BRM/CHR2, the ATPase subunit of the large switch/sucrose non-fermentable (SWI/SNF) complex, has dual roles in transcription of miRNA genes and in inhibiting miRNA processing by remodelling pri-miRNA transcripts associated with SE (58). In addition to the five proteins that are core components of or can associate with the Arabidopsis Microprocessor, six subunits of the THO/TREX complex (THO2, THO1, THO6, THO5A, THO7 and THO4B) co-purified with FLAG-SE. This result demonstrated that the association of SE/ARS2 with the mRNA export machinery is conserved between human and plants (23). Interestingly, ZCCHC8A, a subunit of the Arabidopsis Nuclear Exosome Targeting complex (NEXT) was the most enriched protein in the SE IPs. In fact, all components of the NEXT complex HEN2, ZCCHC8A, ZCCHC8B and RBM7 co-purified with SE (Fig. 1B – red squares). Taken together, our data confirm the previously reported interaction of SE with the nuclear cap binding complex (CBC) and miRNA biogenesis factors, and detect the THO/TREX and NEXT complexes as additional partners of SE in plants.

**Figure 1.**
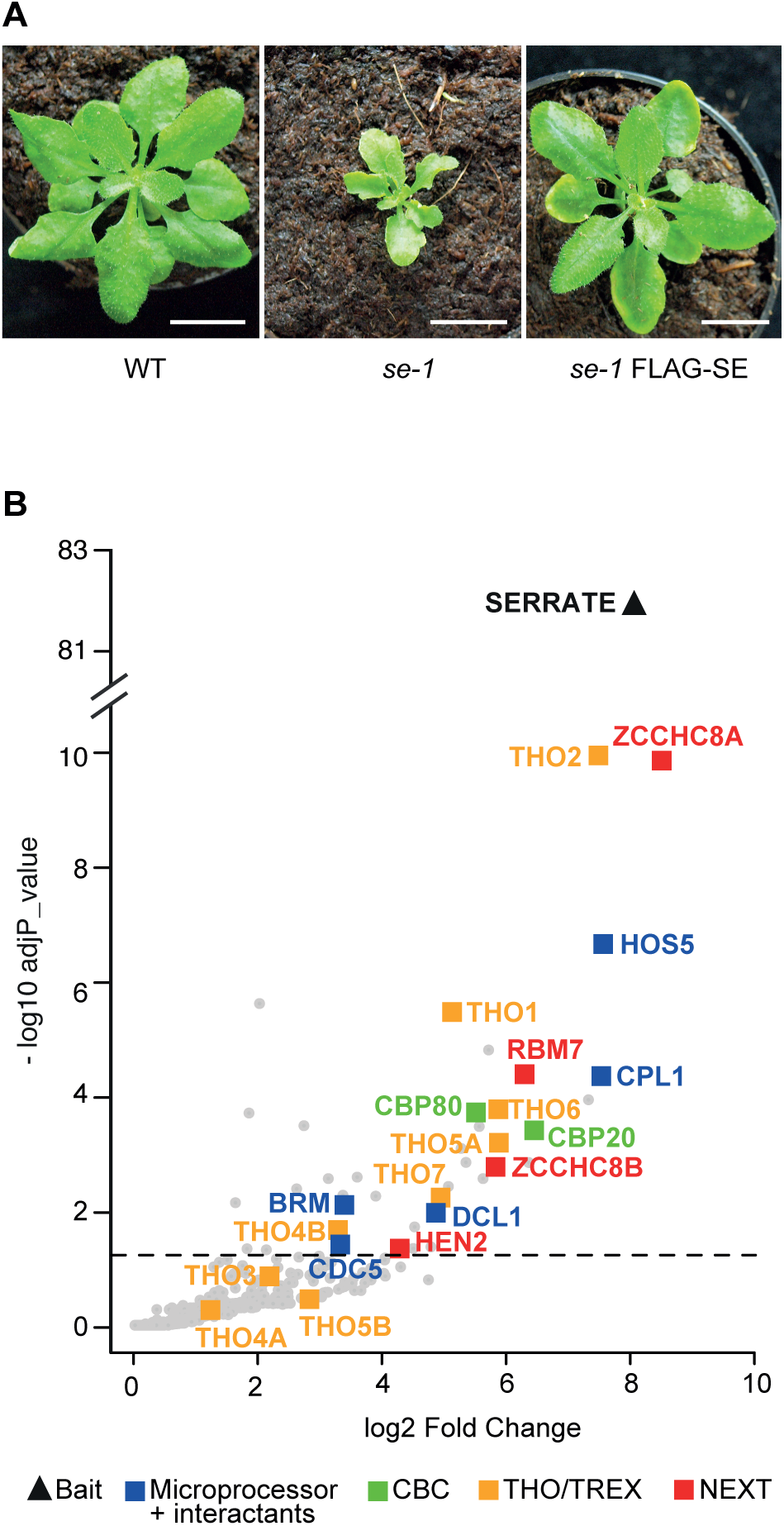
Proteins associated with Arabidopsis SERRATE. **(A)** Phenotypes of 4 week-old wild-type (WT), *se-1*, and *se-1* plants expressing a FLAG-tagged version of SE (FLAG-SE). Scale bars are 1 cm. **(B)** Volcano plot showing the proteins co-purified with FLAG-SE as compared to control IPs performed in Col-0. The bait SE is indicated by a black triangle. Blue squares label the proteins known to compose or interact with the Arabidopsis Microprocessor complex, green squares label the two subunits of the cap-binding complex (CBC), yellow squares label the subunits of the THO/TREX complex, and red squares label the subunits of the NEXT complex. The dashed line indicates the significance threshold (adjp < 0.05).

We then used Arabidopsis lines expressing GFP-tagged versions of ZCCHC8A, ZCCHC8B and RBM7 to test the association between SE and the NEXT complex. Indeed, SE was co-purified when ZCCHC8A, ZCCHC8B or RBM7 were used as baits (Fig. 2, Supplementary Table 3). This result confirmed the association of SE with the NEXT complex. The presence of both ZCCHC8A and ZCCHC8B in SE and RBM7 IPs also confirmed that Arabidopsis NEXT can contain either ZCCHC8A or ZCCHC8B. However, the observation that ZCCHC8A is more enriched than ZCCHC8B indicates that the majority of Arabidopsis NEXT complexes comprises the ZCCHC8A subunit, consistent with the lower expression levels of ZCCHC8B.

**Figure 2.**
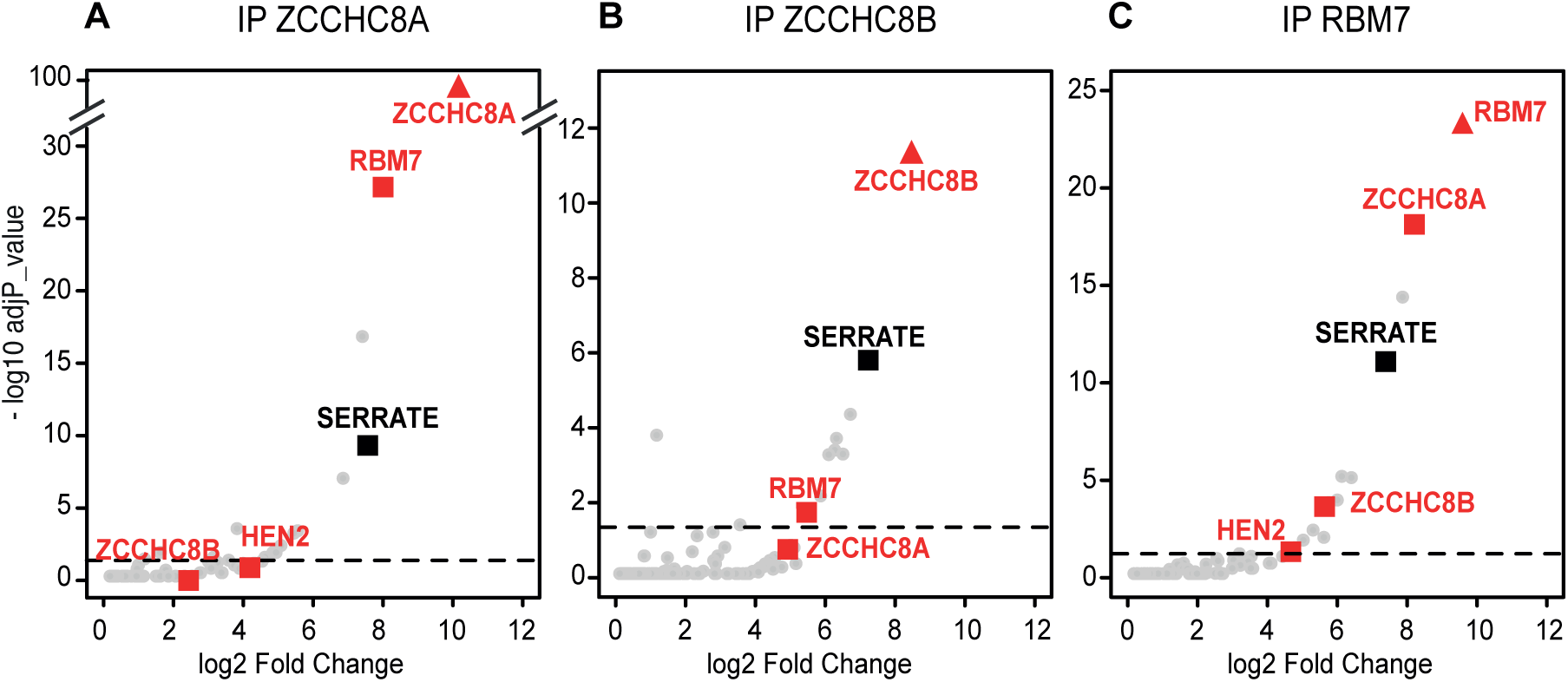
The interaction of SE with the NEXT complex is confirmed by reciprocal IPs. Volcano plots showing the proteins co-purified using GFP-tagged versions of **(A)** ZCCHC8A, **(B)** ZCCHC8B, and **(C)** RBM7 as baits as compared to control IPs performed in Col-0. The bait proteins are indicated by triangles, SE is indicated by a black square, the red squares label the subunits of the NEXT complex. The proteins above the threshold (adjp < 0.05, dashed line) are significantly enriched.

### SE directly contacts ZCCHC8A and ZCCHC8B but not HEN2

As compared to ZCCHC8A and RBM7, the RNA helicase HEN2 was poorly enriched in SE-IPs (Fig. 1B), and previous reciprocal IP experiments using HEN2 as bait (59) did also not detect SE. A possible interpretation of these data is that SE does not directly interact with HEN2 in Arabidopsis. To identify the NEXT subunits that interact directly with SERRATE we first performed yeast two-hybrid (Y2H) assays. SE was fused to the Gal4 DNA binding domain and co-expressed with HEN2, ZCCHC8A, ZCCHC8B or RBM7 fused to the Gal4 activation domain. Only strains co-expressing SE with ZCCHC8A, ZCCHC8B and RBM7 grew under conditions selective for interacting proteins, suggesting that SE can bind to ZCCHC8A, ZCCHC8B and RBM7 (Fig. 3A). By contrast, no interaction was observed between SE and the RNA helicase HEN2 (Fig. 3A). We then tested the interactions between SE and NEXT components in plant protoplasts using Forster resonance energy transfer (FRET) analysed by fluorescence lifetime imaging microscopy (FLIM). The FLIM-FRET method allows to visualize and to quantify direct protein interactions by observing the decrease of the fluorescence lifetime of a donor (SE-GFP in this study) in the presence of an acceptor protein (ZCCHC8A-RFP, ZCCHC8B-RFP, RBM7-RFP, HEN2-RFP in this study). PRP39A-RFP (11) was used as negative control. All fluorescent fusion proteins were expressed to similar levels and co-localised in the nucleoplasm of transformed protoplasts (Supplementary Fig. S1). The fluorescence lifetime of SE-GFP was reduced upon co-expression of ZCCHC8A-RFP, ZCCHC8B-RFP and RBM7-RFP but was unchanged upon co-expression of HEN2-RFP (Fig. 3B). These results were in line with the results obtained by Y2H and support the conclusion that SE directly binds to the ZCCHC8 and RBM7 subunits of NEXT but not to the RNA helicase HEN2. Finally, we also tested the interactions between SE and the NEXT subunits by *in vitro* pull-down assays (Supplementary Fig. S2). Recombinant SE fused to the maltose binding protein (MBP-SE) was used as bait while HEN2, ZCCHC8A, ZCCHC8B and RBM7 proteins were produced as radiolabelled proteins by *in vitro* translation. All samples were treated with RNase to ensure that only direct protein-protein interactions were detected. These experiments confirmed the direct interactions between SE and both ZCCHC8A and ZCCHC8B. By contrast, neither HEN2 nor RBM7 was pulled-down by MBP-SE (Supplementary Fig. S2). The reason for the lack of interaction between SE and RBM7 *in vitro* is unknown.

**Figure 3.**
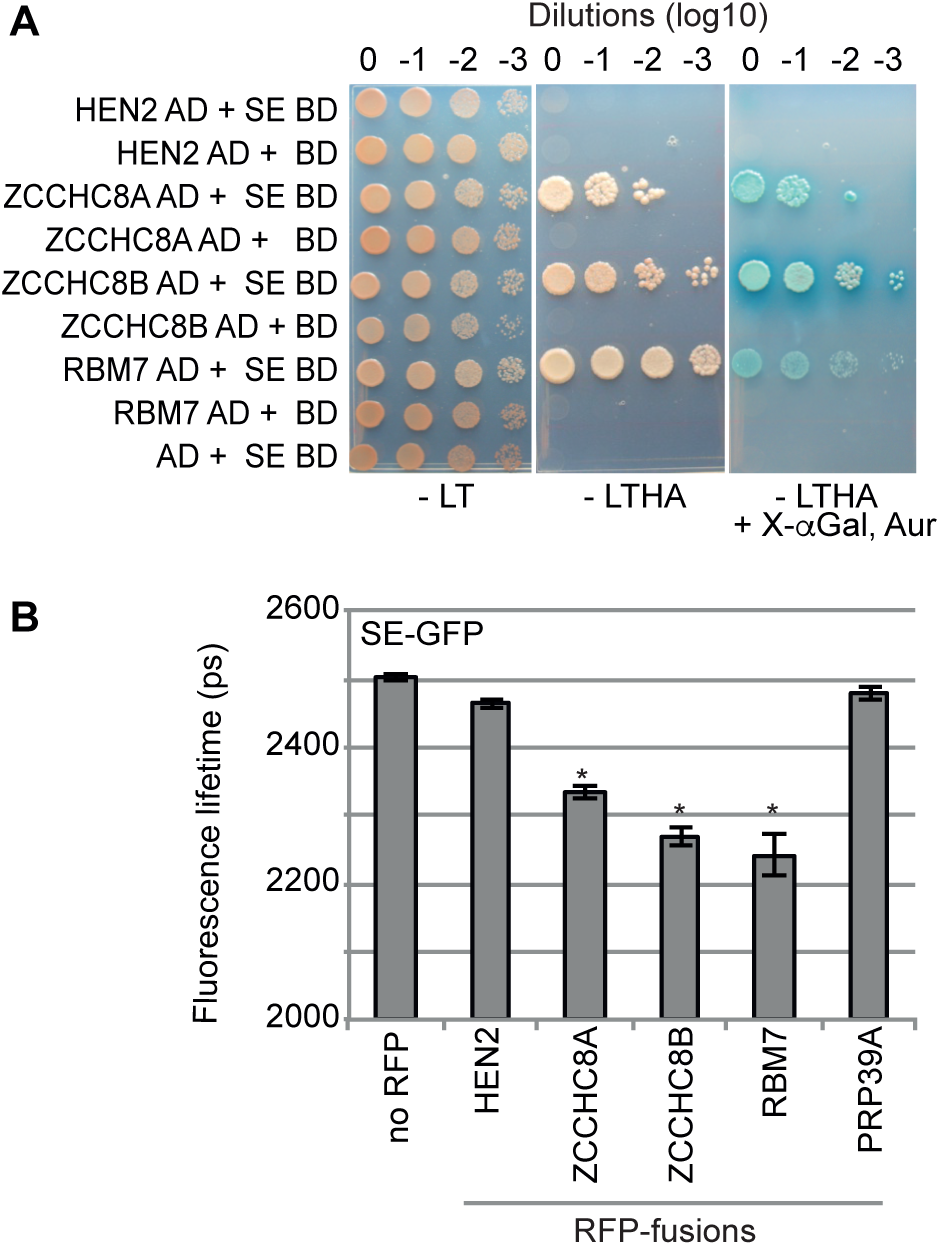
SERRATE interacts with the RBM7 and ZCCHC8 subunits of the NEXT complex. **(A)** Yeast two hybrid assays. SE was fused to the Gal4 DNA binding domain (BD). HEN2, ZCCHC8A, ZCCHC8B and RBM7 were fused to the Gal4 activation domain (AD). Yeast co-expressing the indicated proteins and SE-BD or harbouring the empty pGBDKT7 vector were selected on medium lacking the amino acids leucine (L) and tryptophan (T). Interaction was tested on medium lacking also histidine (H) and adenine (A), as well as on medium complemented with aureobasidin (Aur) and the galactosidase substrate X-αGal. **(B)** FRET-FLIM analyses of Arabidopsis protoplasts expressing GFP fused to SERRATE as fluorescent donor protein. HEN2, ZCCHC8A, ZCCHC8B, RBM7 and PRP39A (as negative control) fused to RFP were used as acceptor proteins. The bar graph presents the fluorescence lifetime of SE-GFP in picoseconds (ps). Error bars indicate the standard error of the mean, n > 10. The asterisks indicate significant differences (p < 0.001, Mann-Whitney-Wilcoxon test) between the control samples co-expressing SE-GFP and PRP39A-RFP, and the samples co-expressing SE-GFP and the indicated RFP fusion proteins.

Both Y2H and FRET-FLIM assays were also applied to test the individual interactions among the subunits of the NEXT complex. These experiments revealed that HEN2 directly interacts with the two zinc finger-containing proteins ZCCHC8A and ZCCHC8B, but not with the RNA binding protein RBM7 (Fig. 4A and B). The interactions between RBM7 and ZCCHC8 proteins could not be tested by Y2H assays because their fusion to the Gal4 DNA binding domain (BD) auto-activated the expression of the reporter gene (Fig. 4A). However, FRET-FLIM analysis indicated that both ZCCHC8A and ZCCHC8B can interact with RBM7-RFP (Fig. 4B). Moreover, FRET-FLIM data obtained with ZCCHC8A and ZCCHC8B as either donors or acceptors suggested that the two proteins can interact and form both homo- and heterodimers (Fig. 4B). To test whether HEN2, ZCCHC8A and SE can form a trimeric complex we combined bimolecular fluorescence complementation (BiFC) and FRET-FLIM. The fluorescent entity formed by the interaction of nVEN-HEN2 with ZCCHC8A-cCFP (Supplementary Fig. S1) was used as a donor in FRET-FLIM with SE-RFP as acceptor. Indeed, the fluorescence lifetime of the BiFC-produced donor formed by nVEN-HEN2-ZCCHC8A-cCFP interaction was reduced upon co-expression of SE-RFP, while the co-expression of the PRP39A-RFP control had no effect (Fig. 4C).

**Figure 4.**
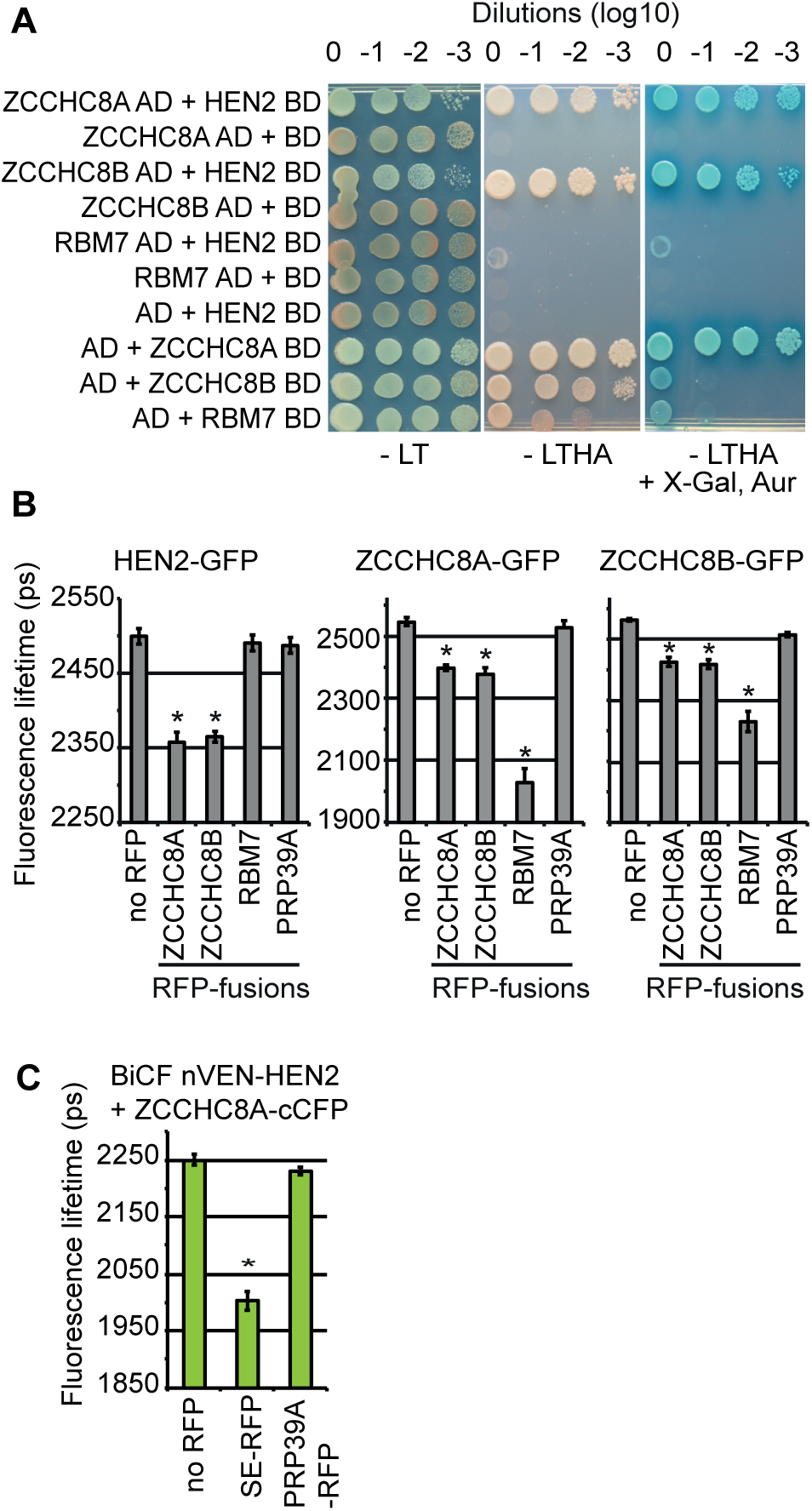
ZCCHC8A and ZCCHC8B interact with both RBM7 and HEN2. **(A)** Yeast two hybrid analyses of the interactions between the NEXT subunits. Medium lacking leucine (L) and tryptophan (T) was used to select for pGADKT7 and pGBDKT7 vectors. Interaction was tested on medium lacking also histidine (H) and adenine (A), and by adding aureobasidin (Aur) and the galactosidase substrate X-αGal. The interactions within the NEXT subunits were evaluated using HEN2-BD as the fusion of ZCCHC8A, ZCCHC8B and RBM7 to the binding domain resulted in autoactivation. **(B)** FRET-FLIM analyses of protoplasts co-expressing fluorescent fusion proteins. The bar graphs present the fluorescence lifetime of the donor molecules HEN2-GFP (left panel), ZCCHC8A-GFP (middle panel) and ZCCHC8B-GFP (right panel) in picoseconds (ps). The RFP-fusion proteins used as fluorescence acceptors are indicated at the bottom. **(C)** SE, HEN2 and ZCCHC8A form a trimeric complex. Combined BiFC and FRET-FLIM analysis. The fluorescent donor molecules are produced by the bimolecular interaction of HEN2 and ZCCHC8A fused to the N- and C-terminal fragments of Venus and Cyan fluorescent proteins, respectively. SE and PRP39A were fused to RFP and served as acceptors. The bar graphs present the fluorescence lifetime of the donor molecule produced by BiFC between nVEN-HEN2 and ZCCHC8A-cCFP in picoseconds (ps). Error bars indicate the standard error of the mean, n > 10. The asterisks indicate significant differences (p < 0.001, Mann-Whitney-Wilcoxon test) between the control samples co-expressing the donor fused to GFP and the indicated acceptors fused to RFP to the samples expressing the donor and the PRP39A-RFP control.

Taken together, our results indicate that the RNA helicase HEN2 and the RNA binding protein RBM7 bind to the two Zn-knuckle proteins ZCCHC8A and ZCCHC8B, but do not bind to each other. In contrast to its human ortholog ARS2 which directly binds to hMTR4 (23), Arabidopsis SE does not bind to HEN2 but can directly interact with each of ZCCHC8A, ZCCHC8B and RBM7.

### The NEXT complex contributes to the degradation of miRNA precursors

Because SE’s substrates notably include miRNA precursors and HEN2 is a key component of the NEXT complex promoting RNA degradation by the nucleoplasmic RNA exosome, we tested for an eventual contribution of HEN2 on the accumulation of miRNA precursors. To this end, RNA-seq libraries were produced and analysed for wild-type and *hen2-2* plants. We focused on the 53 Arabidopsis *miRNA* genes *(MIRs)* with fully annotated gene structures (*i.e.* including transcription initiation and termination sites) that are expressed from independent transcription units, *i.e.* not embedded in introns of protein coding genes. Of those 53 pri-miRNAs, 44 were detected in at least 3/6 libraries and had more reads in *hen2-2* mutants as compared to wild-type plants (Fig. 5A). The analysis of selected loci revealed complex RNA-seq profiles indicating that *hen2* mutant plants accumulate a mixture of miRNA precursors and processing intermediates including introns and maturation by-products corresponding to the regions 5′ and/or 3′ of the mature miRNA sequences (Fig. 5B-H, Supplementary Fig. S3). For several *MIR* genes including *MIR159a*, *MIR159b*, *MIR161*, *MIR172b* and *MIR823*, the RNA-seq distribution profiles suggested also an accumulation of the unspliced and unprocessed primary miRNA transcripts (pri-miRNA) (Fig. 5, Supplementary Fig. S3).

**Figure 5.**
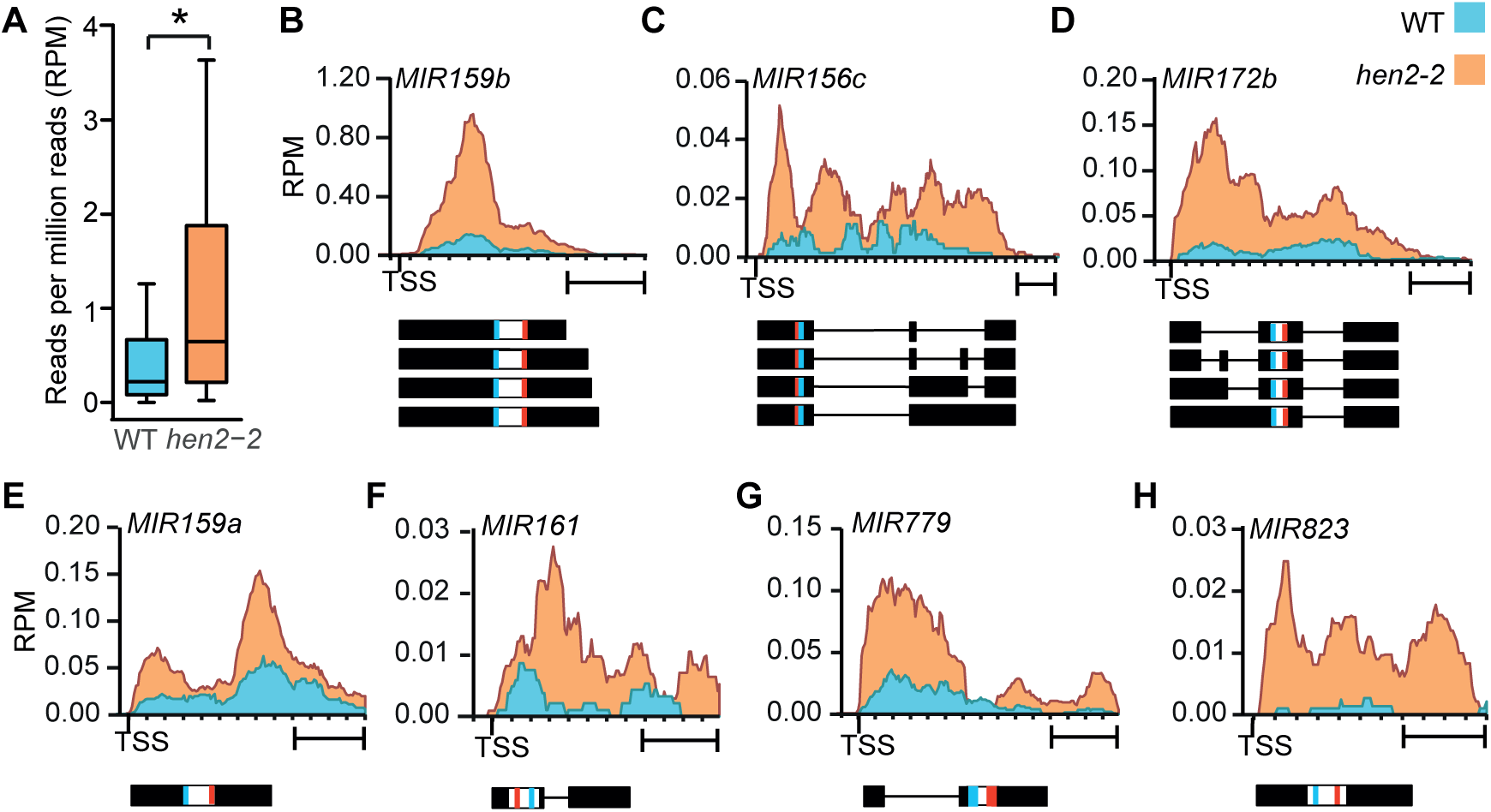
Loss of HEN2 results in the accumulation of miRNA precursors. **(A)** Box plot showing the levels of pri-miRNA transcripts from 44 fully annotated *MIR* genes in 3 biological replicates of wild type (WT) and *hen2-2.* The star indicates the significant difference between the two samples (p < 0.0001, Wilcoxon Signed Rank test) **(B-H)** Coverage of RNA-seq reads shown as reads per million reads (RPM) for selected miRNA genes in WT (blue) and *hen2-2* (orange). The organisation of the *MIR* genes is show below each diagram. Lines represent introns, boxes indicate exons. The miRNA and miRNA* sequences are indicated in red and blue, respectively. TSS, transcription start site. Scale bars are 400 bp. Additional examples are shown in Supplementary Fig. S3.

To test this possibility, we selected two miRNA loci and performed qRT-PCR analysis with primers located in the 5′ region of pri-miRNA that would detect both 5′ maturation by-products and miRNA precursors (Fig. 6A). Fragments corresponding to 5′ maturation by-products were also detectable by 3′ RACE-PCR (Supplementary Fig. S4). A second primer pair was designed to specifically detect pri-miRNAs (i.e. unspliced and/or spliced, but in any case miRNA precursors not processed by DCL1) (Fig. 6A). As compared to wild type, the signals obtained with the primers located in the 5′-regions (blue bar graphs) were 25-fold (5′ pri-miRNA 159a) and 35-fold (5′ pri-miRNA 161) higher in *hen2-2* plants. The signals obtained with the primers specific to the pri-miRNAs (red bar-graphs) were 10-fold (pri-miRNA 159a) and 8-fold (pri-miRNA 161) higher in *hen2-2* mutant plants. These results confirmed the results obtained by RNA sequencing and support the conclusion that *hen2-2* mutants accumulate pri-miRNAs, in addition to the 5′ maturation by-products which were also observed in previous studies (35,60). The same experiment was also performed in mutants of NEXT subunits. While *zcchc8a-1* and *zcchc8b-1* single mutants had levels similar to wild type, *zcchc8a-1 zcchc8b-1* double mutants also accumulated pri-miRNAs 159a and 161, suggesting that ZCCHC8A and ZCCHC8B can substitute for each other. Loss of the RNA binding protein RBM7 also resulted in the accumulation of pri-miRNAs (Fig. 6A).

**Figure 6.**
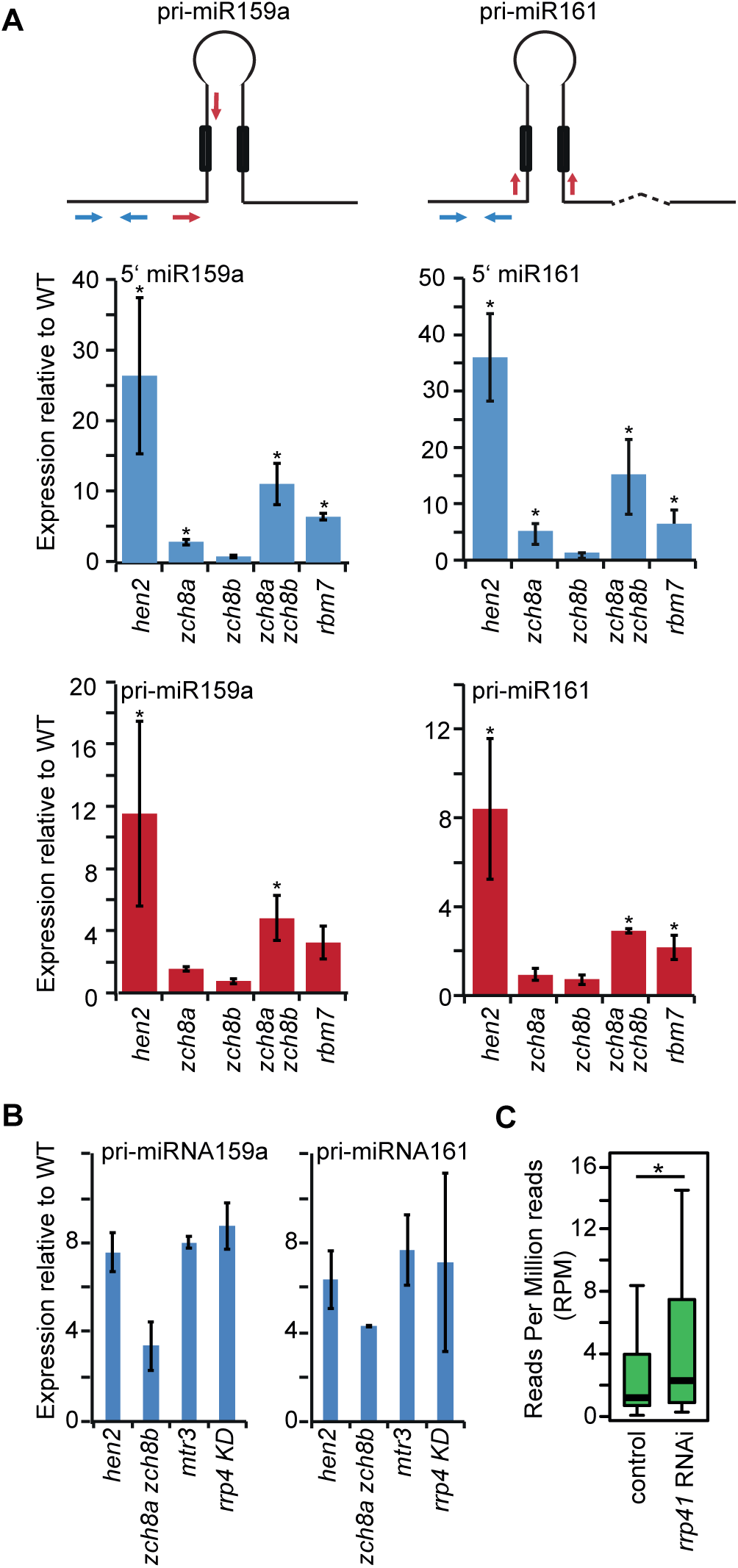
Accumulation of maturation by-products and miRNA precursor transcripts in NEXT mutants. **(A)** Quantification of pri-miRNA levels by qRT-PCR in mutants of NEXT components. The diagrams shown at the top illustrate the primary transcripts of pri-miRNAs 159a and 161. The dashed line indicates an intron. The boxes indicate the miRNA and miRNA* sequences within the stem-loop. The first dicing step occurs at the edges of the stem-loop and releases both 5‘ and 3 maturation by-products and the pre-miRNA which is diced again to excise the mature miRNA/mRNA* duplex. The primers specific to the 5‘ regions of pri-miRNAs are indicated by blue arrows, the red arrows indicate the primers that detect the pri-miRNA. The bar graphs show the mean of three biological replicates, the error bar represents the SD, asterisks indicate significant accumulation (p < 0.05, Student’s t-test) as compared to wild type. **(B)** Quantification of pri-miRNA levels by qRT-PCR in exosome mutants. Bar graphs show the mean of two replicates, the error bars indicate the SD. **(C)** Boxplot showing the global levels of 42 miRNA precursors with known gene structures in *rrp41* RNAi and control samples. The RNA-seq data for this analysis were retrieved from (61). The star indicates a significant difference between the means (Wilcoxon Signed Rank Test, p< 0.05).

Because the NEXT complex activates RNA degradation by the RNA exosome, we then determined whether these pri-miRNA precursors would also accumulate in RNA exosome mutants. Indeed, we also detected accumulation of pri-miRNAs 159a and 161 in *mtr3* and *rrp4 KD*, two viable mutants of the exosome core complex (Fig. 6B). Similar results were obtained for additional pri-miRNA transcripts that we tested by qRT-PCR (Supplementary Fig. S5). Finally, we re-analysed a recently published RNA-seq dataset (61) obtained from *rrp41* RNAi mutants for the accumulation of all pri-miRNAs with known gene structure. 42 *MIR* transcripts were detected in at least half of the libraries and showed an overall accumulation in *rrp41* RNAi mutants as compared to controls (Fig. 6C), strongly indicating that pri-miRNAs are indeed substrates of the exosome complex.

Taken together, these data support the conclusion that NEXT contributes to the degradation of miRNA maturation by-products and assists in the elimination of surplus pri-miRNAs by the RNA exosome.

### Common substrates of HEN2 and SE include targets of nuclear RNA surveillance

The phenotypic analysis of *hen2-2*, *se-2* and *hen2-2 se-2* plants provided additional genetic evidence of a functional link between HEN2 and SE. While *hen2-2* plants have no obvious growth defects, *se-2* mutants display a pleiotropic developmental phenotype with hyponastic leaves characteristic for many miRNA biogenesis mutants (10). In addition, the margins of *se-2* mutant leaves are serrated. Interestingly, the simultaneous mutation of *HEN2* and *SE* partially restored the leaf phenotypes displayed by single *se-2* mutants (Fig. 7), although *se-2 hen2-2* plants are sterile when grown under standard conditions. To investigate the molecular basis for the partial restoration of wild-type phenotypes in *se-2 hen2-2*, we first determined the transcriptomes of wild-type, *hen2-2*, *se-2* and *se-2 hen2-2* plants by RNA-seq. Each genotype was compared to wild type using DESeq2 (Fig. 8, Supplementary Fig. S6, Supplementary Table 4). Down-regulated genes were not further analysed because of the very limited overlap among the three genotypes (Supplementary Fig. S6B). Of the 544 genes upregulated in *hen2-2*, 411 were also upregulated in *se-2 hen2-2* (Fig. 8A, Supplementary Fig. S6C). In agreement with the previously reported roles of HEN2 in RNA surveillance, the 263 genes that were specifically affected in *hen2-2* and the double mutants but remained unchanged in *se-2* comprised a large proportion of long non-coding RNAs (22.2 %), snoRNAs (10.8 %), snRNAs (1.4 %) and other non-coding RNAs (1.4 %) which likely represent direct targets of HEN2-mediated RNA degradation (Fig 8B, Supplementary Table 4). Of the 590 genes upregulated in *se-2*, 369 were also detected in *se-2 hen2-2* (Fig. 8A, Supplementary Fig. S6C). The 211 genes that were specifically upregulated in *se-2* and *se-2 hen2-2* but not affected in *hen2-2* were mainly miRNA genes (8.1 %) and protein coding genes (88.2 %) (Fig. 8B, Supplementary Table 4). The common transcripts, upregulated in *hen2-2*, *se-2* and *se-2 hen2-2*, comprised protein coding genes (73 %), *MIR* genes (10.8 %), long non-coding (lnc) RNA (6.8 %), ncRNA (2 %), snRNA (1.4 %) and snoRNA transcripts (6.1 %) (Fig. 8B, Supplementary Fig. 7). The majority of the common transcripts (131/148) showed a similar accumulation in *se-2 hen2* as compared to single mutants, in line with the idea that SE and HEN2 act together to promote their degradation. By contrast, 265 of the transcripts that were upregulated in *se-2 hen2-2* plants showed significantly increased levels in *se-2 hen2-2* as compared to each of the single mutants (Fig. 8C, Supplementary Table 4). Interestingly, this group of transcripts was mainly composed of snRNAs (Fig. 8D), protein coding genes which often showed a particular increase in reads mapping to the 5′ region and the first intron (Fig. 8E), and miRNA precursors (Fig. 8F).

**Figure 7.**
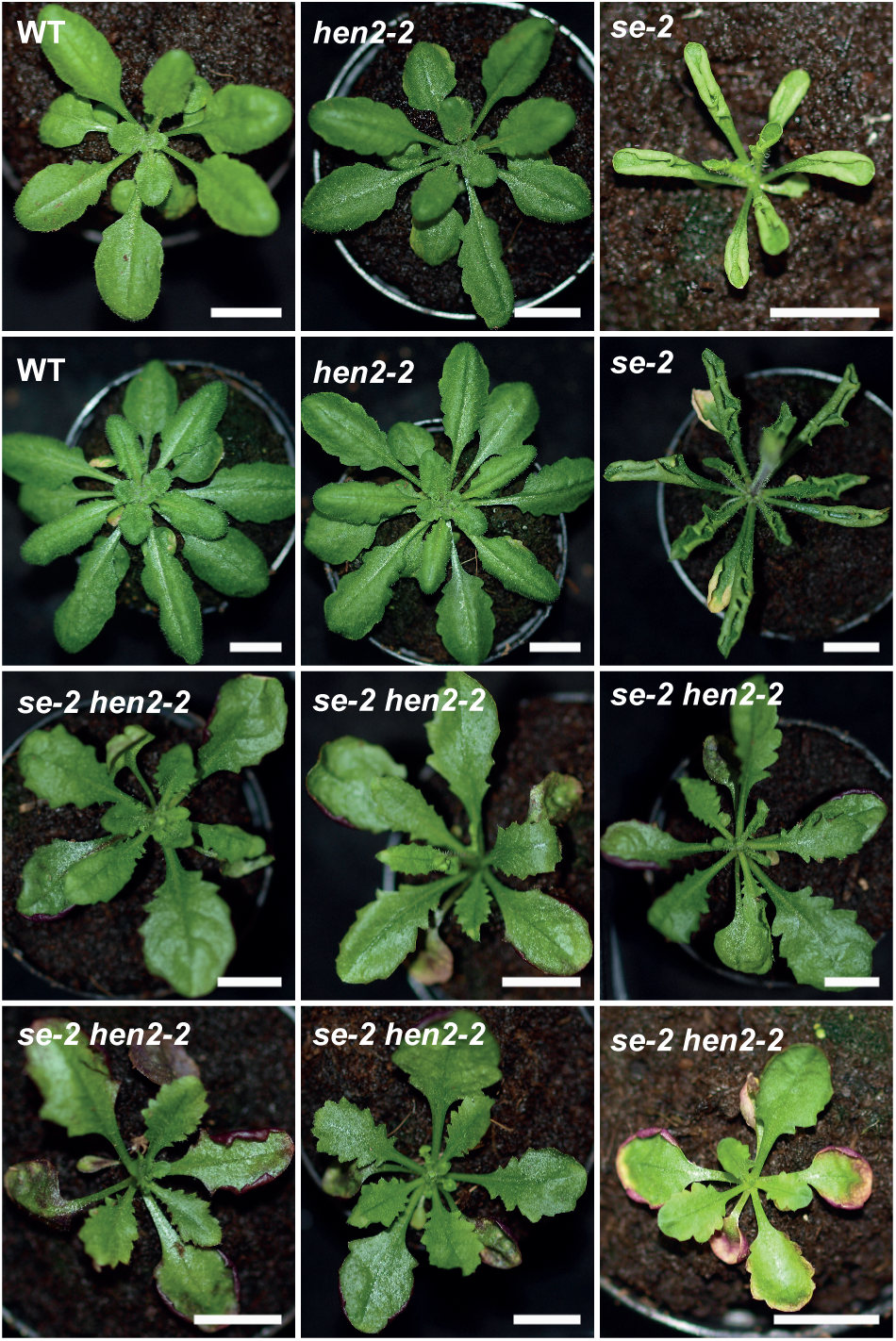
Mutating HEN2 partially restores the phenotypes of *se-2* plants. 5 week-old plants of the genotypes indicated in each panel. Scale bars are 1 cm.

**Figure 8.**
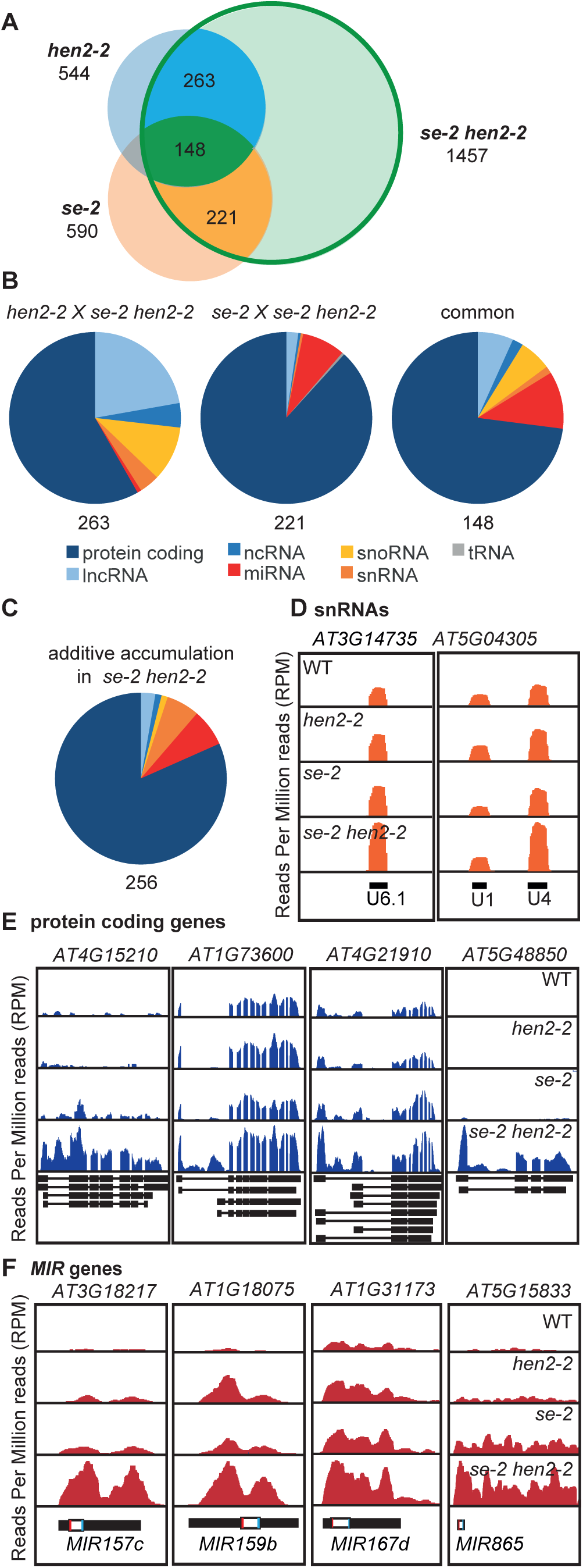
Common targets of both SE and HEN2. **(A)** Genome-wide transcriptome analysis by RNA-seq. Each genotype was compared to WT by DESeq2. The Venn diagram displays the numbers of upregulated transcripts in *hen2-2, se-2* and *se-2 hen2-2* samples. **(B)** Pie charts displaying the proportions of different types of transcripts upregulated in both *hen2-2* and *se-2 hen2-2* (left, intense blue area in (A)), *se-2* and *se-2 hen2-2* (middle, intense yellow area in (A)) and in all three genotypes (right, intense green area in (A)). Dark blue, protein coding genes; light blue, long non-coding RNA genes; royal blue, other non-coding RNA genes; red, miRNA genes; yellow, snoRNA genes; orange, snRNA genes; grey, tRNA genes. **(C)** Pie chart showing the transcript types that accumulate to higher levels in *se-2 hen2-2* as compared to each of the single mutants. (**D-F)** RNA-seq read distribution profiles for representative loci encoding snRNAs (D), protein coding genes with increased reads in their 5′ regions, (E), and *MIR* genes (F). The diagrams at the bottom of each panel show the location and structure of the annotated genes with boxes as exons and lines as introns. White boxes represent pre-miRNA, red and blue bars represent mature miRNA and miRNA*, respectively.

### Compromised pri-miRNA degradation compensates for poor miRNA production

43 of the 53 pri-miRNAs with known gene structure were detected in at least 6/12 libraries and had overall more reads in *se-2 hen2-2* samples as compared to *hen2-2* or *se-2* samples (Fig. 9A, Supplementary Table 5). A similar trend was observed for miRNA precursors that are incompletely annotated (Supplementary Fig. S8). To investigate the impact of pri-miRNA accumulation on the levels of mature miRNAs, we analysed wild-type, *hen2-2*, *se-2* and *se-2 hen2-2* plants by small RNA-seq. This experiment revealed that loss of HEN2 had no global effect on the accumulation of mature miRNAs, while more that 120 miRNAs had significantly reduced levels in *se-2* mutants (Fig. 9B, Supplementary Fig. S9). In wild type and *hen2-2* samples, 23 and 20 % of small RNAs corresponded to annotated miRNAs, respectively. By contrast, only 5.8 % of the total reads were mature miRNAs in *se-2* (Fig. 9B). Mutating HEN2 in the *se-2* background partially restored global miRNA levels to 9.4 %. We determined the fold changes for 53 miRNAs with known gene structure and their corresponding miRNAs by DESeq2. Interestingly, we detected a correlation between the increase of pri-miRNAs and the increase of the corresponding mature miRNAs for the comparison of *se-2 hen2-2* versus *se-2*, but not for any of the other pairwise comparisons (Fig. 9C, Supplementary Fig. S10). This result supports the idea that pri-miRNA levels are not limiting in wild type, *hen2-2 and se-2*, but are linked to restored miRNA levels in *se-2 hen2-2*. The restoration of miRNA function is also reflected by the fact that many of the miRNA target mRNAs that over-accumulate in *se-2* as consequence of compromised miRNA function have close to wild-type levels in *se-2 hen2-2* (Fig. 9D, Supplementary Table 4).

**Figure 9.**
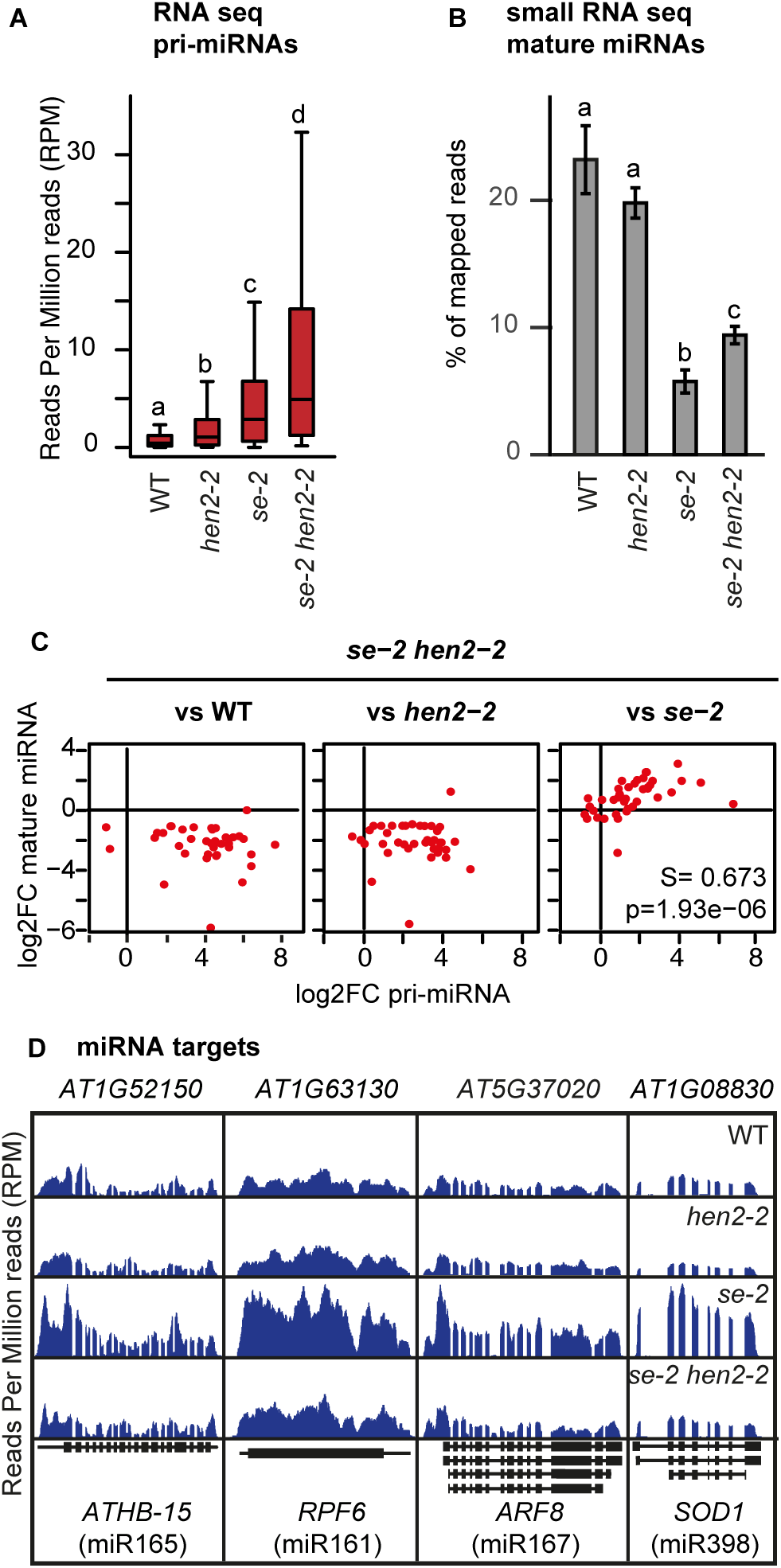
Accumulation of pri-miRNA precursors partially restores miRNA production in *se-2 hen2-2* double mutants. **(A)** Boxplot showing the levels of 43 primary miRNA precursors with known gene structures that were detected in at least 6 of 12 RNA-seq libraries. Different letters indicate significant differences (p < 0.0001, Wilcoxon Signed Rank test) **(B)** Bar graph showing the levels of mature miRNAs determined by small RNA-seq in 3 biological replicates of each genotype in percent of total mapped reads. Different letters indicate significant difference between the samples (p < 0.05, Student’s t-test). **(C)** RNA-seq and small RNA-seq data were compared by DESeq2 and the fold-change of the 43 primary miRNAs considered in (A) was plotted against the fold-change of the corresponding mature miRNAs. A significant positive correlation was determined when *se-2 hen2-2* was compared to *se-2*, but not for any of the other comparisons. S, Spearman correlation coefficient; p, p-value. **(D)** RNA-seq read distribution profiles showing the accumulation of known miRNA target mRNA in *se-2* and the restoration of near to wild-type levels in *se-2 hen2-2*. The diagrams below each panel illustrate the annotated transcripts with boxes for exons and lines for introns.

Interestingly, mutating HEN2 also partially rescued the developmental defects of *hyl1-2* mutants (Fig. 10A). Unlike mutating SE, loss of HYL1 had no effect on the accumulation of some known targets of HEN2 (Supplementary Fig. 11), but *hyl1-2* mutant plants accumulate miRNA precursors transcripts due to impaired miRNA maturation (3,6). qRT-PCR analysis demonstrated that *hyl1-2 hen2-2* double mutants also accumulated higher levels of pri-miRNAs 159a, 161 and 165 as compared to single *hen2-2* or *hyl1-2* mutants (Fig. 10B). Finally, small RNA blots revealed that *hyl1-2 hen2-2* double mutants accumulated also more mature miRNAs 159 and 161 than *hyl1-2*, similar to what we observed in *se-2 hen2-2* (Fig 10C). These data further support the hypothesis that higher levels of miRNA precursors resulting from impaired pri-miRNA degradation can compensate for compromised miRNA production efficiency.

**Figure 10.**
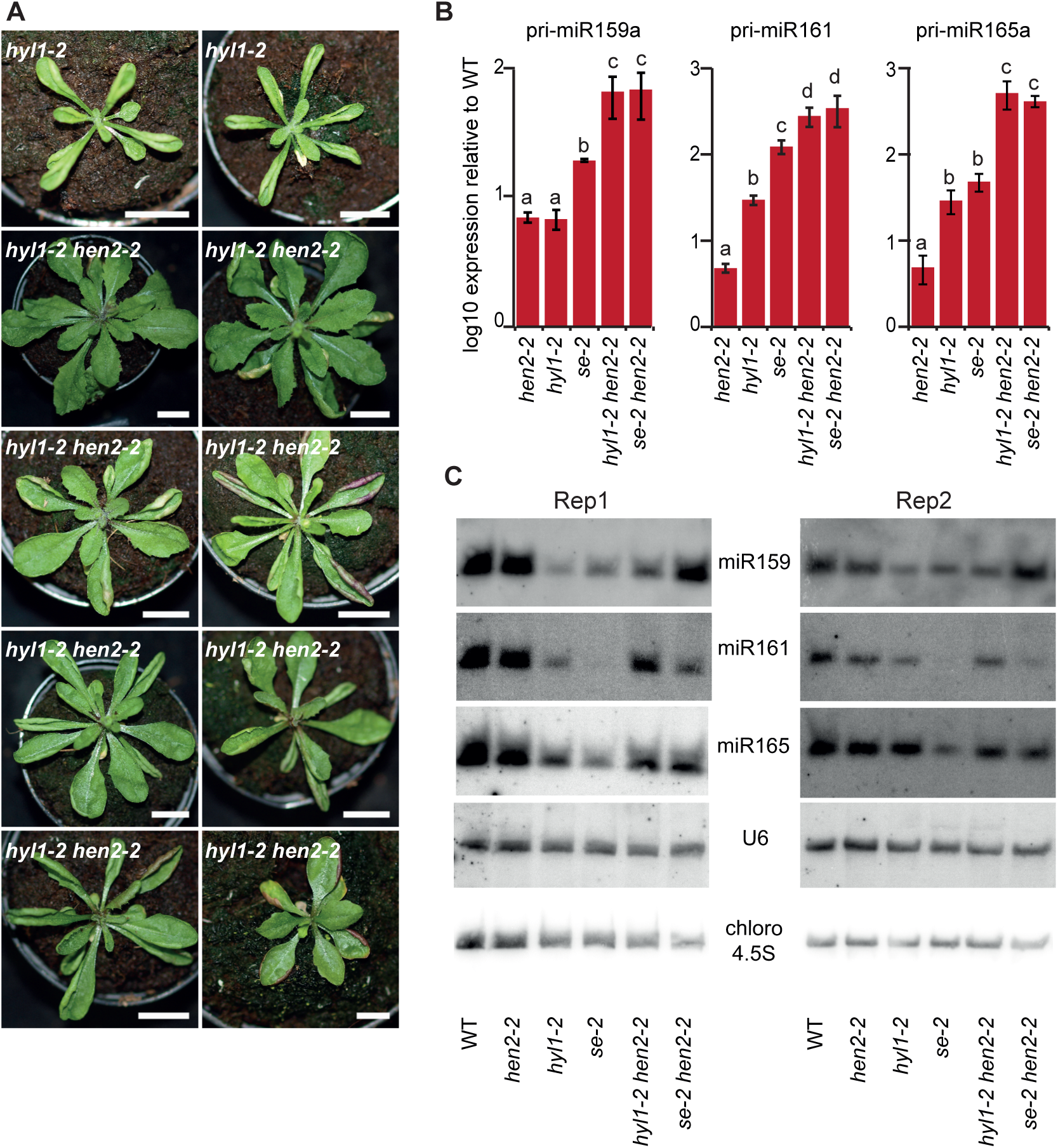
Simultaneous loss of HYL1 and HEN2 partially rescues decreased miRNA. **(A)** Representative pictures of 4 week-old *hyl1-2* and *hyl1-2 hen2-2* plants. Scale bars are 1 cm. **(B)** Relative expression levels of three miRNA precursors determined by qRT-PCR in three biological replicates of wild type, *hen2-2*, *hyl1-2*, *se-2*, *hyl1-2 hen2-2* and *se-2 hen2-2*. Distinct letters indicate difference between the means (p < 0,05, Student‘s t-test). Error bars show the SD. **(C)** Small RNA blots of two biological replicates (Rep) were hybridised with probes specific to miRNAs and U6 snRNA as indicated. Hybridisation to chloroplast 4.5 ribosomal RNA is shown as a loading control.

## DISCUSSION

A major result of this study is the unbiased identification of proteins associated with Arabidopsis SE. The proteins that co-purified with FLAG-SE can be sorted in four main groups. The first group comprised the two subunits of the cap-binding complex. The direct and stable interaction between CBC and SE/ARS2 is conserved between human and plants and provides the basis for the role of SE/ARS2 as adapter protein between capped RNA complexes and various RNA maturation complexes, including those for 3′ end processing, transport and degradation (15,62). The second group comprises proteins required for different steps in the production of mature miRNA from primary miRNA transcripts. Each of HOS5, CPL1, BRM and CDC5 has previously been shown to physically associate with SE and at least one other component of the trimeric Microprocessor core complex, DCL1 or HYL1 (54,55,57,58). However, only DCL1 was enriched in SE IPs while HYL1 was not detected. This may be linked to technical reasons, perhaps the protein or its association with SE is destabilized under our experimental conditions. Another possible explanation is that the interaction of SE with HOS5 and CPL1 occurs also independently of the Microprocessor complex and promotes transcription and processing of other types of RNAs.

As a third group of SE-interacting proteins we identified homologs of several subunits of the THO/TREX complex. The THO/TREX complex is an evolutionary conserved assembly that connects transcription elongation, splicing and export of mRNAs (63). Arabidopsis THO/TREX is also required for the biogenesis of miRNAs (64,65). The core complex of Arabidopsis THO/TREX consists of HPR1 (THO1), THO2, THO5A/B, THO6, THO7A/B and TEX1 (66) associated with UAP56, MOS11 and the mRNA export adapters THO4B-C (aka ALY2-4) (67). Using TEX1, UAP56 and MOS11 as baits, Arabidopsis THO/TREX was shown to associate with a number of splicing factors and with components of the nuclear pore complex (67). Our data now extend these observations by showing that the association of THO/TREX with SE/ARS2 is also conserved in plants.

Finally, a portion of Arabidopsis SE is associated with the NEXT complex, similarly to its human counterpart ARS2 (21). The association of SE with the NEXT complex was confirmed by multiple reciprocal co-immunoprecipitations. Our data strongly indicate that this association is mediated by the ZCCHC8 subunits, which also directly interact with both the HEN2 and RBM7 subunits within the NEXT complex. This latter observation suggests that ZCCHC8A and ZCCHC8B, similar to the role of hZCCHC8 in human NEXT (54), function as central scaffold proteins of NEXT in Arabidopsis. Interestingly, our FRET-FLIM data indicate also that each of ZCCHC8A and ZCCHC8B can interact with itself and with each other. This opens the possibility that the NEXT complex may contain two (or more) ZCCHC8 subunits, and that heterogenic NEXT complexes containing both ZCCHC8A and ZCCHC8B may also exist. However, as compared to ZCCHC8A, poor enrichment of ZCCHC8B in RBM7 or SE-IPs strongly suggests that the majority of Arabidopsis NEXT contains ZCCHC8A. Whether the incorporation of ZCCHC8A and/or ZCCHC8B influences the functional properties of NEXT remains to be studied in detail. In this study, loss of either ZCCHC8A or ZCCHC8B had no major effect and accumulation of miRNA precursors was only observed in the double *zcchc8a zcchc8b* mutant. Hence, ZCCHC8A and ZCCHC8B have at least partially overlapping functions and can replace each other. Moreover, the duality of ZCCHC8A and ZCCHC8B genes is not systematically conserved in higher plants indicating that most species possess a single NEXT complex similar to metazoa.

An interesting observation is that the molecular interaction between NEXT and SE/ARS2 is differently mediated in human and plants. According to *in vitro* assays, ARS2 can directly interact with hMTR4 (23), the functional homolog of Arabidopsis HEN2 (and its nucleolar counterpart AtMTR4) in human cells. The competition of hMTR4 and THO4 for the association with hARS2 determines whether an RNA molecule is exported to the cytosol or degraded by the nuclear exosome (23). In addition, the interaction of human NEXT and PAXT with the ARS2-CBC complex is facilitated by the Zn-finger protein ZC3H18. In our experiments, none of the proteins that were co-purified with SE or the NEXT subunits shared convincing homology with ZC3H18, and the hMTR4 homolog HEN2 was rather poorly enriched in SE IPs and did not directly interact with SE in any of our assays. By contrast, our data strongly indicate that Arabidopsis SE recruits the NEXT complex via a direct interaction with the ZCCHC8 subunits. How these variations are linked to the differences in protein structure (24) and how this impacts RNA processing and stability will be an interesting focus of future research. It also remains to be investigated whether ZCCHC8A/B competes with components of the Arabidopsis THO/TREX complex for binding to SE in a similar manner as hMTR4 and human THO compete for binding to ARS2. It is also tempting to speculate that SE binding to NEXT may perhaps exclude the simultaneous binding of miRNA processing factors such as HYL1.

Based on the results presented here we propose that SE participates in RNA degradation through the recruitment of the NEXT complex. It is therefore interesting to note that the double mutant *se-2 hen2-2* is sterile while double *hyl1-2 hen2-2* show normal fertility. Whether the sterility of *se-*2 *hen2-2* is linked to the role of SE in RNA degradation which is not shared by HYL1 is unknown to date. However, although being caused by both direct and indirect effects, the massive perturbance of the transcriptome observed in *se-2 hen2-2* highlights the importance of nuclear RNA surveillance when RNA processing and sorting processes, many of which do involve the SE/ARS2-CBC complex (24,34), are disturbed. Indeed, further relevance of the interaction between SE and the NEXT complex can be inferred from the experimental identification of well-known substrates of exosome-mediated RNA surveillance as common targets (35,60). An particularly interesting observation is the increased accumulation of snRNA transcripts in *se-2 hen2-2* double mutants. Our approach cannot discriminate between processing and degradation of snRNA transcripts, but the fact that levels of mature U6 snRNAs are not reduced in *hen2-2*, *se-2* and *se-2 hen2* would be in line with the idea that both HEN2 and SE are involved in the degradation of misprocessed or surplus transcripts rather than in snRNA production. Interestingly, human ARS2 was shown to stimulate the 3′ end processing of snRNAs, thereby preventing readthrough transcription (20). Hence, a reasonable speculation is that SE/ARS2 has a double role in snRNA metabolism: the binding of SE/ARS2 to nascent snRNA transcripts may promote both processing and degradation, depending on SE’s interaction with either CPL1, triggering 3′ cleavage, or NEXT, inducing degradation by the exosome (19,24,33,34).

Another type of transcripts that we and other teams observed in *hen2-2* and *se-2* plants are truncated or misprocessed transcripts spanning the first intron (14,15,35,68). Similar 5′ transcript derived from protein coding genes have also been observed in human cells depleted from ARS2 and result from premature transcription termination rather than from compromised splicing of full-length pre-mRNAs (19,34,69). However, both SE and ARS2 do interact with splicing factors. Among the proteins that co-purify with FLAG-SE, both HOS5 and CPL1 have been implicated in splicing (56), and a previous study detected several auxiliary factors of the U1 snRNP as SE interactants (70). A very recent study suggests that the ARS2-mediated recruitment of U1 snRNP to 5′ proximal pre-mRNA sequences favours splicing and full-length transcription, and disfavours premature transcription and degradation (69). Hence, the increased accumulation of truncated or mis-spliced transcripts in *se-2 hen2-2* double mutants may also be explained by a dual role of SE in recruiting either splicing factors or NEXT.

Previous studies based on tiling microarrays already observed the accumulation of 5′ fragments excised from miRNA precursors upon downregulation of the RNA exosome or loss of HEN2 (35,60). Our data extend the role of 3′-5′ degradation by the RNA exosome in miRNA metabolism by showing that loss of HEN2, the central co-factor of the nucleoplasmic RNA exosome, results also in the accumulation of pri-miRNAs. The increase in miRNA precursors in *hen2*, *rbm7* and *zcchc8a zcchc8b* mutants indicated that NEXT participates to the degradation of primary miRNA transcripts that are transcribed in excess. This conclusion is further supported by the observation that mutating HEN2 in *se-2* and *hyl1-2* plants, which have higher levels of precursors because dicing is slow and inefficient when SE or HYL1 are absent (3,4), has additive effects and leads to a pronounced accumulation of pri-miRNAs. These conclusions are in line with a growing body of evidence obtained in yeast, human and plants indicating that RNA degradation by the nuclear exosome does not only eliminate misprocessed and defective RNAs but also controls the levels of functional RNA molecules by restricting the pool of precursors available for maturation (71–73). We have previously shown that slowing down the degradation of unspliced pre-mRNA compensates for the inefficient splicing of a splice-site mutation in the *pas2-2* mutant (74). The competition between degradation and processing pathways is also a compelling explanation for the partial rescue of mature miRNA levels observed in *hyl1-2 hen2-2* and *se-2 hen2-2* mutants.

The observation that miRNA precursors accumulate to higher levels in both s*e-2 hen2-2* and *hyl1-2 hen2-2* as compared with their respective single mutants can be explained with the contribution of HEN2 to pri-miRNA degradation. However, the interaction of SE with NEXT and the observation that *se-2* mutants accumulate also typical targets of nuclear RNA-surveillance supports the hypothesis that the SE’s binding to pri-miRNAs can promote both, miRNA processing by recruiting the Microprocessor complex, or RNA degradation triggered by the recruitment of the exosome. Therefore, we propose that SE has a dual role in both miRNA biogenesis and pri-miRNA degradation.

## ACKNOWLEDGEMENTS

Authors would like to thank Dr. Agata Stepien for her help in FRET-FLIM analyses and Dr. Hélène Zuber for her assistance in analyses of mass spectrometry data.

