## Supplementary Figures S1-S11 for "SERRATE interacts with the nuclear exosome targeting (NEXT) complex to degrade primary miRNA precursors in Arabidopsis"

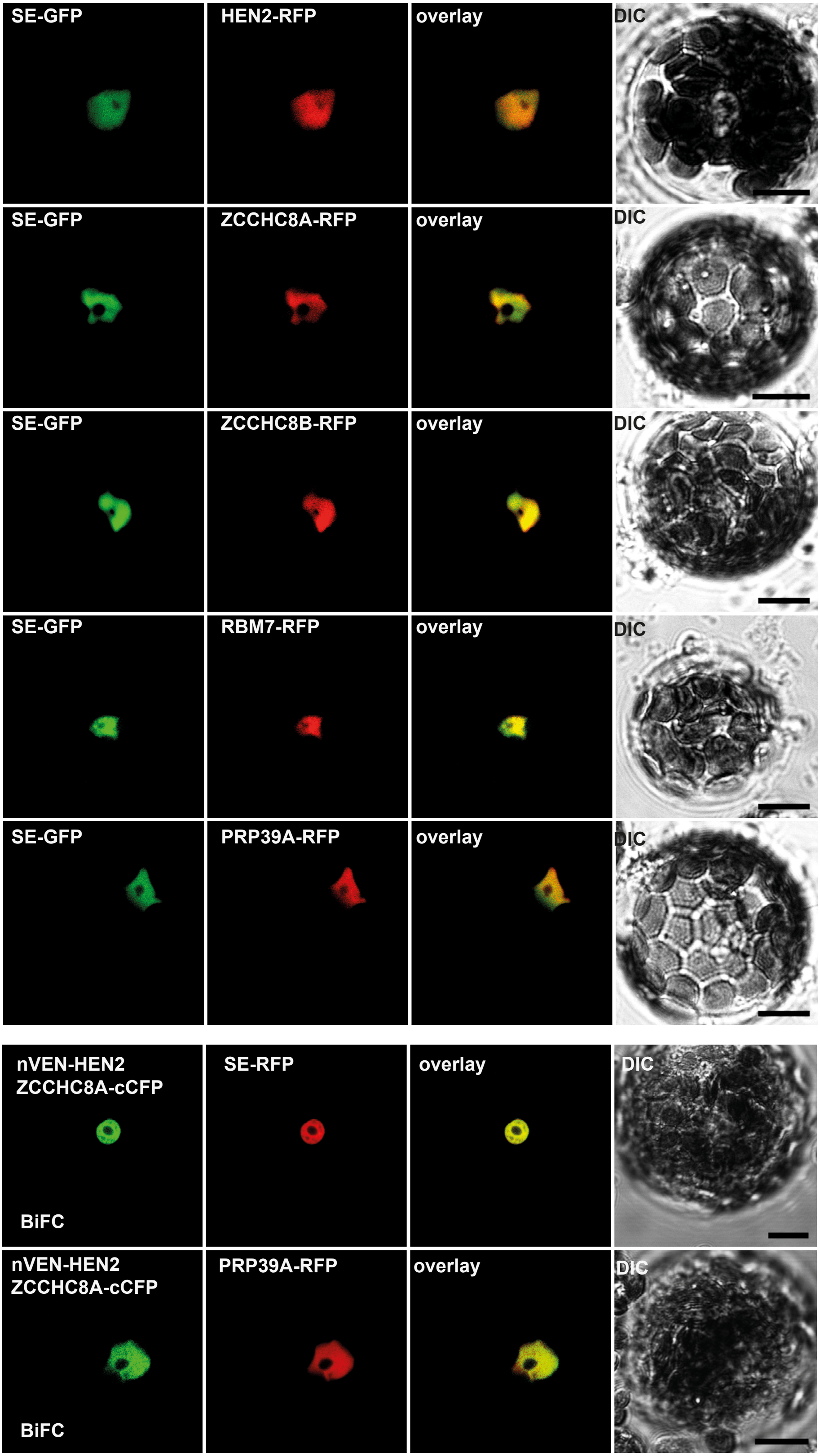


**Supplementary Fig. S1. SE-GFP co-localises with NEXT subunits in the nucleus.** Transient co-expression of fluorescent proteins in Arabidopsis protoplasts. BiFC, Bimolecular Fluorescence Complementation; DIC, differential interference contrast. Scale bars are 10 μm.

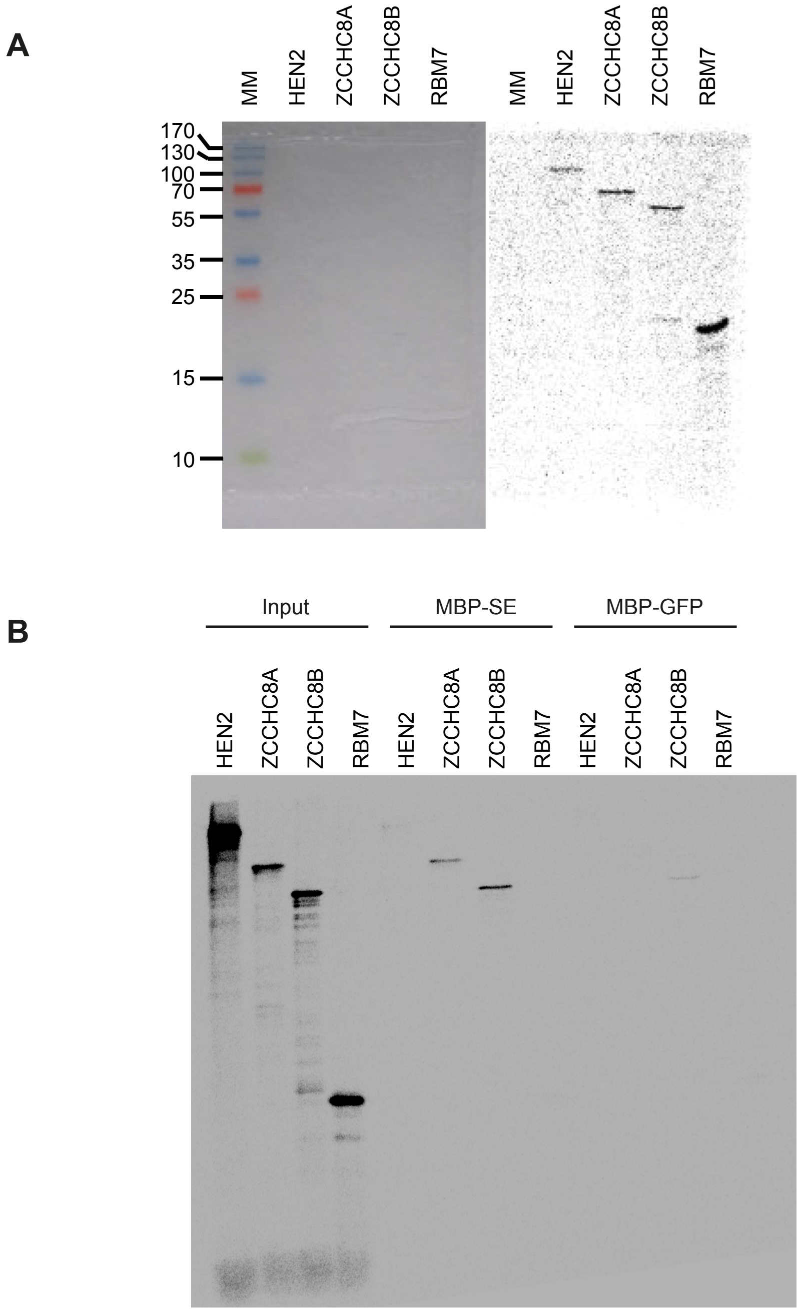


**Supplementary Fig. S2. Recombinant SE interacts with ZCCHC8A and ZCCHC8B.**
**(A)** The proteins indicated on the top of the gel were produced by *in vitro* transcription and translation in presence of [$^{35}$S]-methionine and analysed by autoradiography. The gel with the size marker (in kDa) is shown on the left, the exposure to a phosphoimager plate is shown on the right. (**B)** SE or GFP fused to the maltose binding protein (MBP) were purified from *E. coli,* bound to amylose beads and incubated with radiolabelled proteins as indicated. Proteins were eluted with maltose, separated by SDS-PAGE and visualized using a phosphoimager.

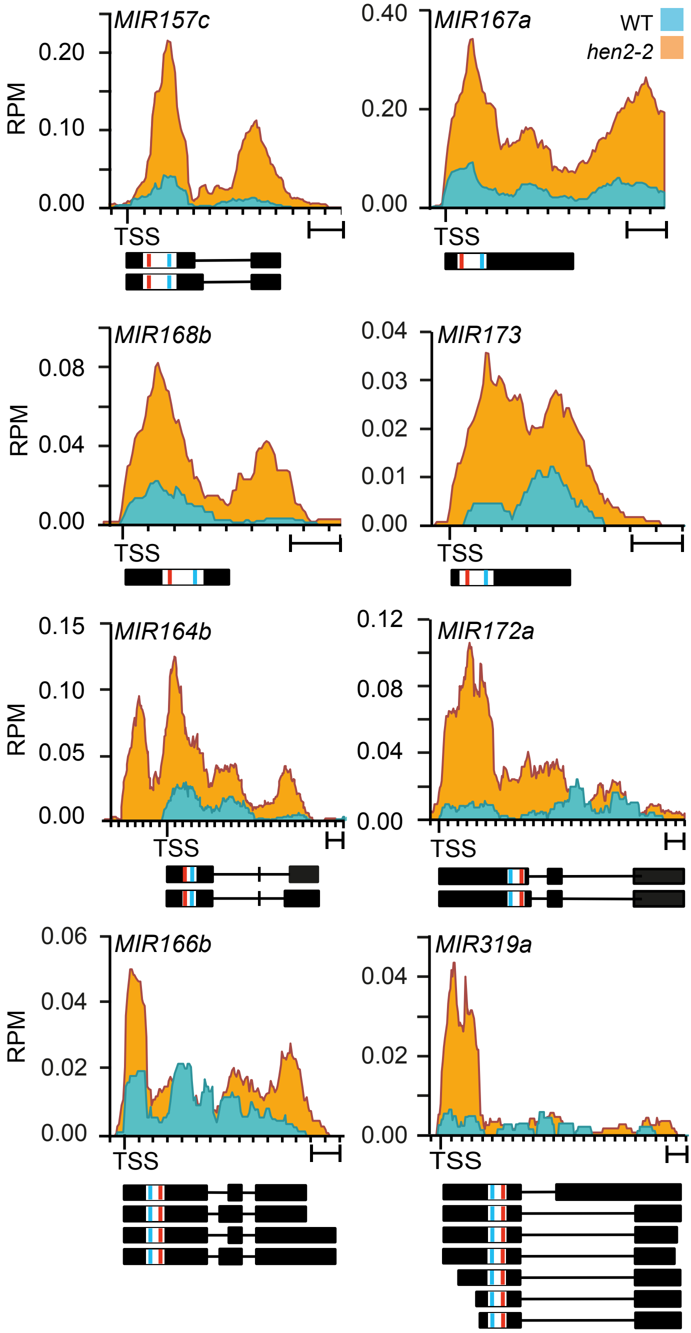


**Supplementary Fig. S3. Accumulation of miRNA precursors and maturation by-products in *hen2-2* plants**. Reads per million reads (RPM) for selected miRNA genes in WT (blue) and *hen2-2* (orange). The organisation of the miRNA genes is show below each diagram. Lines represent introns, boxes indicate exons. White boxes correspond to pre-miRNAs. The location of mature miRNA and miRNA* sequences are indicated in red and blue, respectively. TSS, transcription start site. Scale bars are 200 bp.

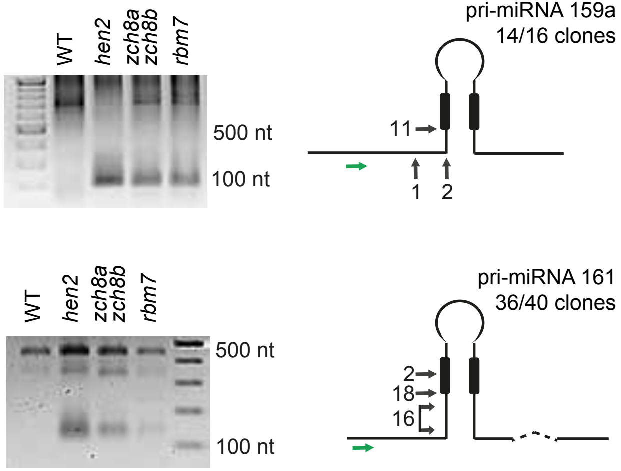


**Supplementary Fig. S4. 5' by-products of miRNA processing detected by 3‘ RACE.** Following cDNA synthesis primed with oligo-dT, 3' RACE-PCR with a primer (green arrow) situated in the 5' region of the pri-miRNA sequence was used to detect 5' maturation by-products. PCR products were analysed by electrophoresis (left). The PCR products detected in *hen2-2* mutants were excised, cloned and sequenced (right). A diagram of the pri-miRNA is shown on the right. The black boxes indicate miRNA and miRNA* sequences. The dashed line indicates an intron. The forward primer used for 3‘ RACE is indicated by the green arrow. The black arrows indicate the 3' ends of the cloned PCR products.

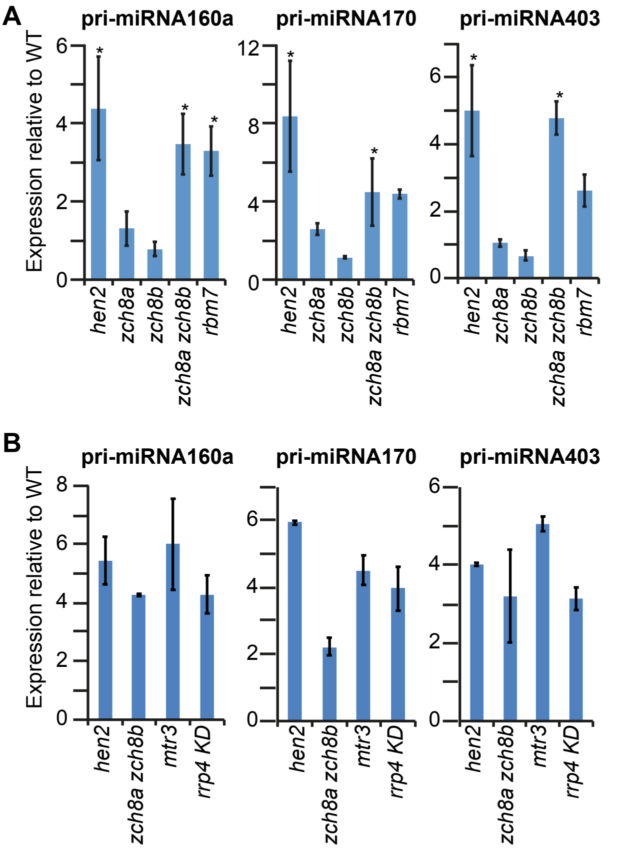


**Supplementary Fig. S5. Accumulation of pri-miRNAs in NEXT and exosome mutants.** Levels of pri-miRNAs were determined by qRT-PCR using primers specifically detecting pri-miRNAs. The bar graphs show the mean of three (A) and two (B) biological replicates. Asterisks indicate significant accumulation ($p < 0,05$, Student‘s t-test) as compared to wild type.

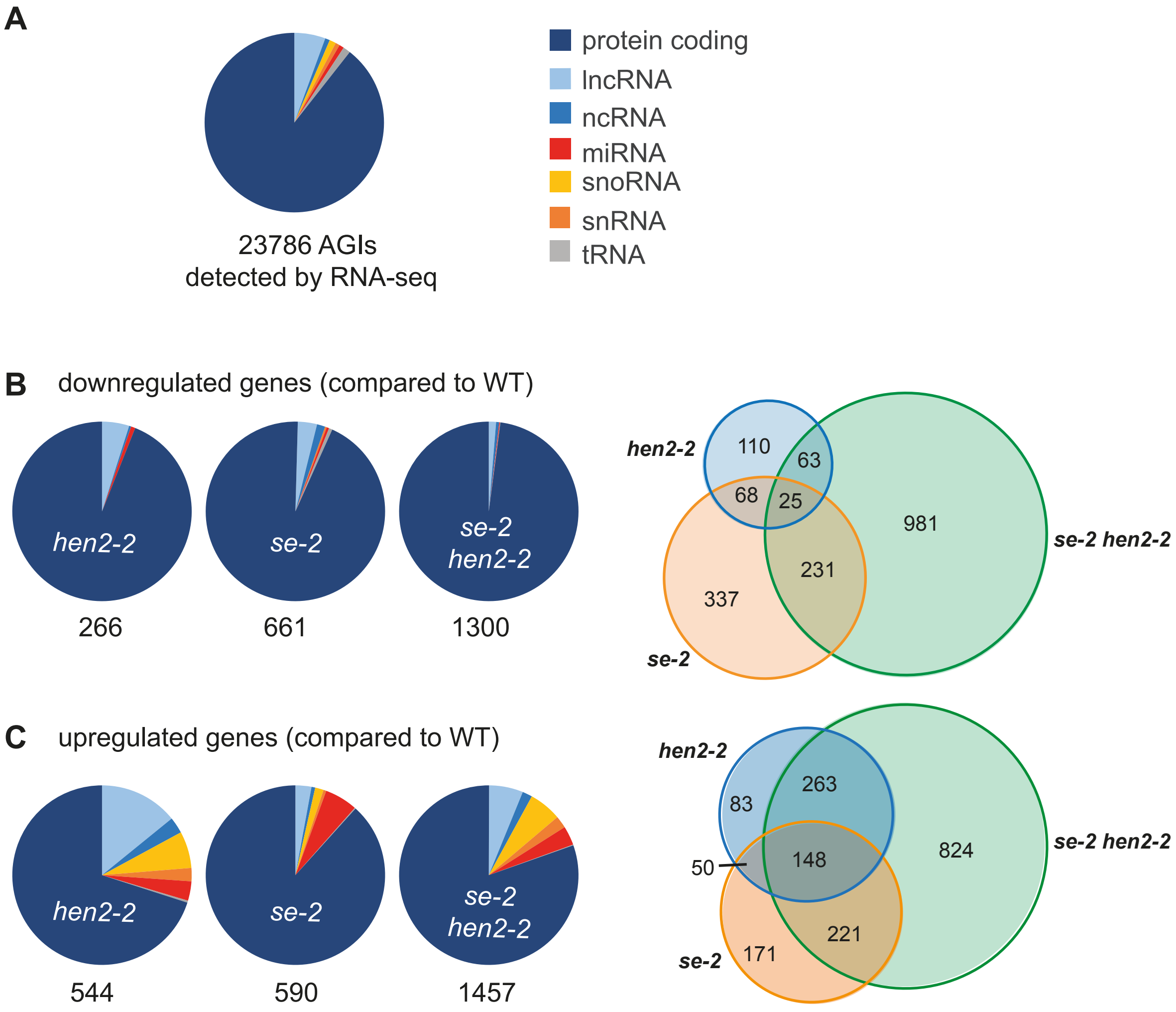


**Supplementary Fig. S6. Comparison of the transcriptomes of wild-type and mutant plants by RNA-seq. (A)** Pie chart showing the proportions of different types of RNAs among the 23786 annotated transcripts that were detected by RNA-seq. **(B, C)** DESeq2 analyses. Each of the RNA-seq datasets obtained from *hen2-2*, *se-2* and *se-2 hen2-2* samples was compared to wild type. The pie charts show the proportion of different types of RNAs that were significantly downregulated (B) or upregulated (C) in each genotype. The numbers below the charts indicate the total number of down- or upregulated genes. The Venn diagrams on the right display the numbers of specific and common transcripts for both downregulated (B) and upregulated genes (C).

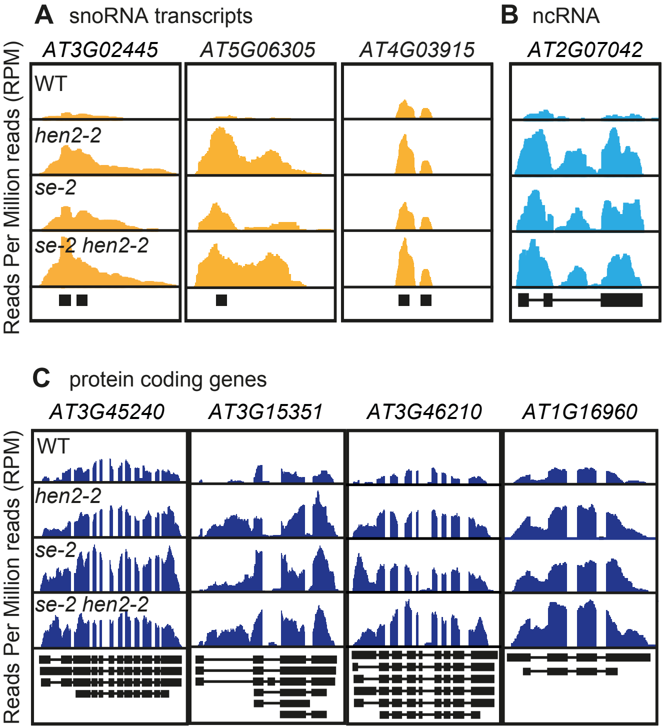


**Supplementary Fig. S7. Both *se-2* and *hen2-2* mutants accumulate substrates of nuclear RNA surveillance.** Common transcripts detected in each of *hen2-2*, *se-2* and *se-2 hen2-2*. Read distribution profiles of representative loci encoding snoRNAs (A), ncRNA (B) and protein coding genes with increased number of reads in their 5' regions (C). The schemes below each panel display the annotated genes with boxes as exons and lines as introns.

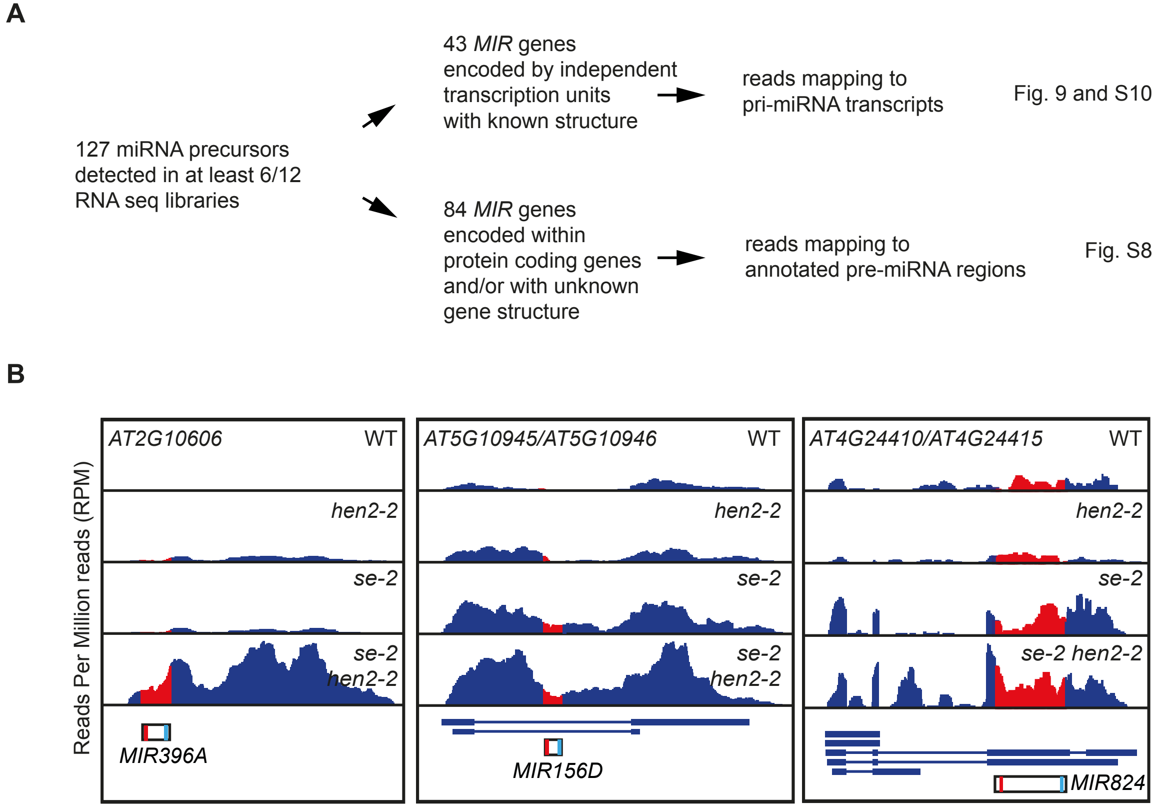


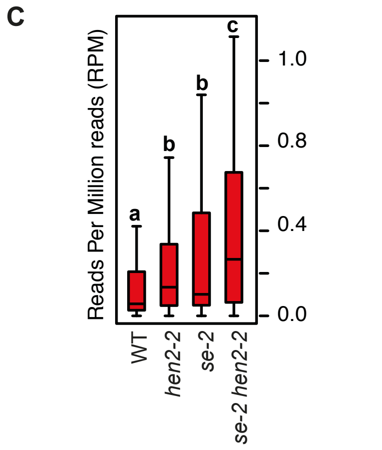


**Supplementary Fig. S8. Detection of primary miRNA precursors by RNA-seq. (A)** Diagram illustrating the analysis of pri-miRNA levels in the second RNA-seq dataset comprising a total number of 12 libraries, three biological replicates for each WT, *hen2-2*, *se-2* and *se-2 hen2-2*. **(B)** RNA-seq read distribution profiles for examples of the 84 miRNA-encoding loci with undetermined gene structure. All regions are shown in 5'-3' orientation of the encoded transcripts. The regions marked in red correspond to the known pre-miRNA sequences. The diagrams below display the annotated genes in these regions. Annotated pre-miRNAs are shown as white boxes, red and light blue bars mark the location of mature miRNAs and miRNA*, respectively. Annotated protein-coding genes are shown in dark blue, with lines for introns and boxes for exons. **(C)** Boxplot calculated from the reads mapping to the pre-miRNA regions of the 84 annotated miRNA genes with unknown gene structure. Different letters mark significant differences between the samples (Wilcoxon Signed Rank test, $p < 0.02$).

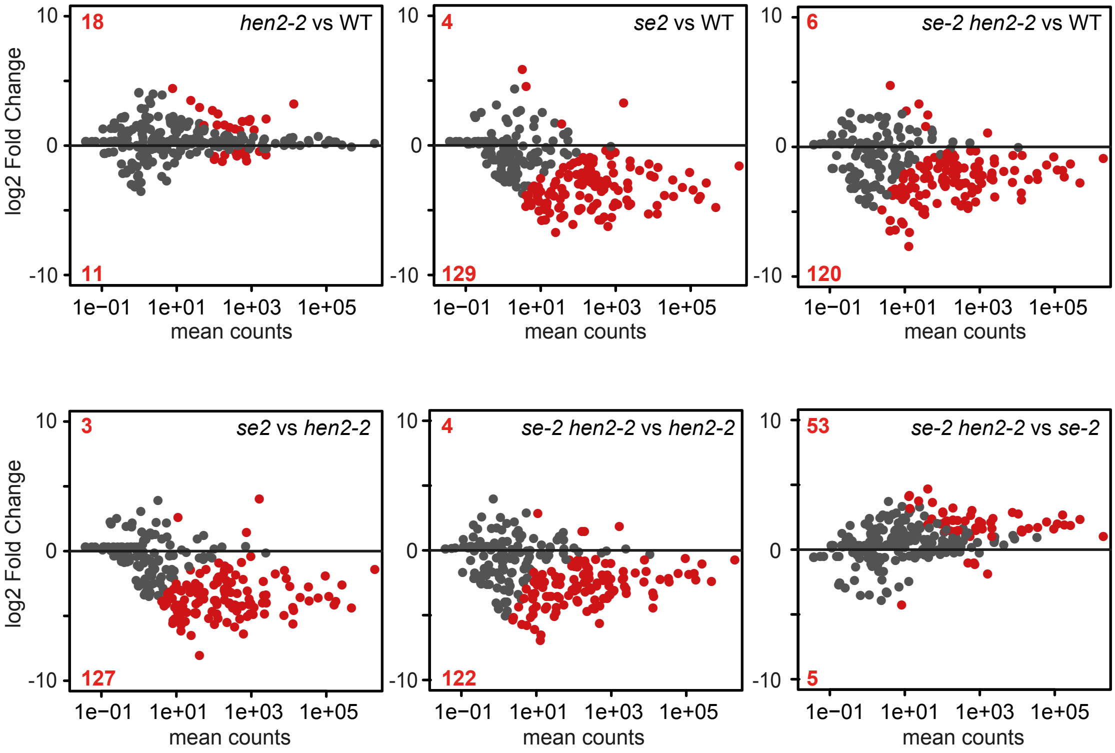


**Supplementary Fig. S9. The *hen2-2* mutation partialy restores levels of mature miRNAs in the *se-2* background.** Pairwise comparisons of small RNA-seq data using DESeq2. For each miRNA, the log2 fold change was plotted against its normalised mean count. miRNAs that showed a significantly different accumulation are labelled in red. The number of miRNAs that are up-or downregulated for each comparison is given in the left corner of each panel.

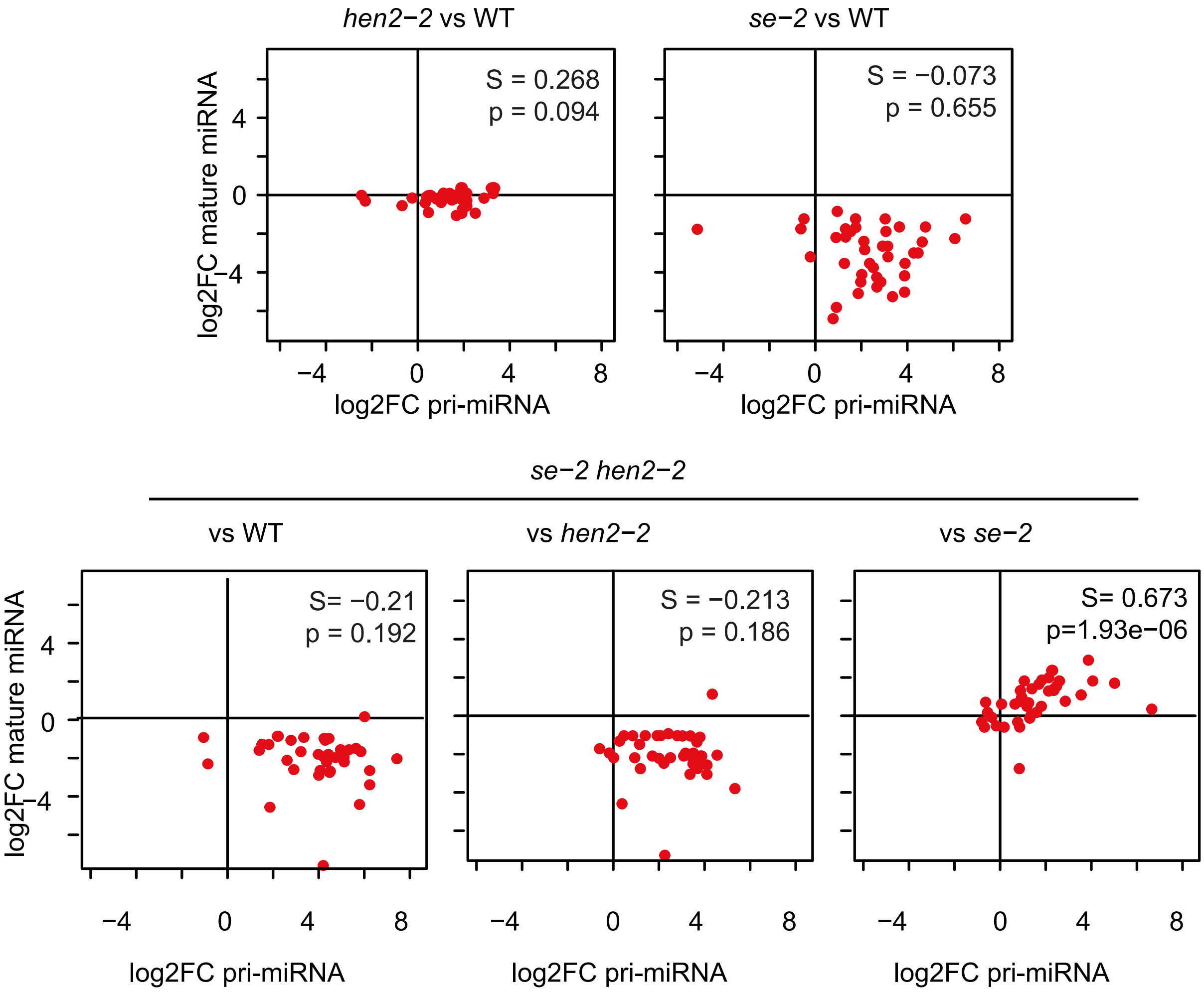


**Supplementary Fig. S10. Restoration of miRNA levels in *se-2 hen2-2* double mutants**. The datasets for both RNA-seq and small RNA-seq data were compared by DESeq2. For each pairwise comparison, the fold-change of the 43 primary miRNAs with known gene structures and detected in 6/12 libraries was plotted against the fold-change of the corresponding mature miRNAs. Note that the three panels at the bottom are also shown in Fig. 9C. S, Spearman correlation coefficient; p, p-value.

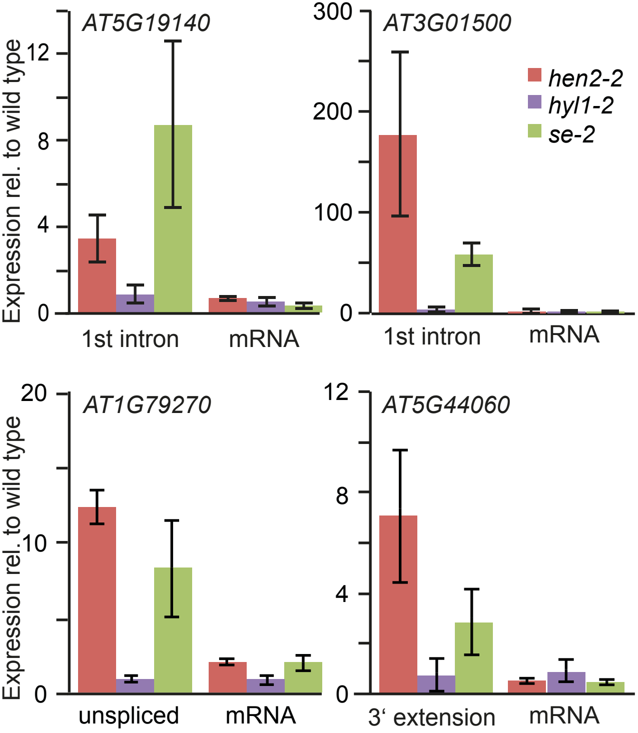


**Supplementary Fig. S11. HYL1 is not required for the degradation of nucleoplasmic exosome targets**. Accumulation of known RNA substrates determined by qRT-PCR in wild-type, *hen2-2*, *hyl1-2* and *se-2* plants. The bar graphs show the relative expression as compared to wild type as mean of three replicates, the error bars indicate the SD.
